# Neural dynamics and architecture of the heading direction circuit in a vertebrate brain

**DOI:** 10.1101/2022.04.27.489672

**Authors:** Luigi Petrucco, Hagar Lavian, You Kure Wu, Fabian Svara, Vilim Štih, Ruben Portugues

## Abstract

Animals can use different strategies to navigate. They may guide their movements by relying on external cues in their environment or, alternatively, by using an internal cognitive map of the space around them and their position within it. An essential part of this representation are heading cells, neurons whose activity depends on the heading direction of the animal. Although those cells have been found in vertebrates, the full network has never been observed and there is very little mechanistic understanding of how these cells acquire their response properties. In this study, we use volumetric functional imaging in larval zebrafish to observe, for the first time in a vertebrate, a full network that encodes allocentric heading direction. This network of approximately one hundred inhibitory neurons is arranged in an anatomical circle in the anterior hindbrain. Its activity is driven purely by the integration of internally generated signals, indicating that a simple vertebrate brain can encode maps of how an animal moves within its surroundings. Single cell reconstructions of electron micrographs allow us to uncover how the connectivity pattern of neurons within the network supports the implementation of a ring attractor network. The neurons we identify share features with neurons in the dorsal tegmentum nucleus of rodents and the fly central complex, showing that similar connectivity and mechanistic principles underlie the generation of cognitive maps of heading direction across the animal kingdom.

---

In many animals, effective navigation in the world involves the use of cognitive maps that provide a representation of position and orientation with respect to the environment. While the position in space has been shown to be encoded in place cells and grid cells in the mammalian hippocampal and entorhinal circuits (Moser et al., 2008), allocentric orientation is represented by head direction cells, neurons that are active any time the animal faces a particular direction in space.

Head direction cells were originally described in the postsubicular cortex (Taube et al., 1990), but have since been observed in several other cortical and subcortical areas (reviewed in (Taube, 2007)). The activity in these head direction networks can be understood in terms of ring attractor networks, where local recurrent excitation is combined with long-range out-of-phase inhibition to create a stable localized bump of activity that encodes direction. This model has received remarkable empirical validation with the observation of heading direction representations in the insect central complex, where key components of a ring attractor network have been mapped onto its neuronal architecture (Seelig and Jayaraman, 2015; Green et al., 2017; Kim et al., 2017; Fisher et al., 2019; Suver et al., 2019; Lyu et al., 2022). However, such mechanistic understanding in vertebrates is still lacking.

The lowest region of the vertebrate brain where head-direction related signals have been found is the dorsal tegmental nucleus (DTN) (Sharp et al., 2001), a paired, GABAergic nucleus located in the brainstem, that originates from rhombomere 1 (Puelles, 2016). In rodents the DTN is closely associated with the interpenducular nucleus (IPN), a hindbrain structure believed to support the DTN heading direction representations (Quina et al., 2017) that has been indirectly implicated in spatial navigation (Clark and Taube, 2009; Clark et al., 2009). In addition, recent studies in larval zebrafish have suggested an important role for the IPN in directional behavior (Dragomir et al., 2020; Cherng et al., 2020). Based on this, we decided to leverage the optical accessibility of the larval zebrafish as a model organism to comprehensively image the anterior hindbrain (aHB) of this vertebrate, in order to identify any potential network activity that could be involved in the encoding of heading direction.

Using a combination of volumetric lightsheet imaging, 2-photon imaging and electron microscopy we uncover a circuit contained within rhombomere 1 that represents heading direction by a persistent and localized bump of activity. This activity profile smoothly translates across the neuronal population as the fish turns, mimicking the compass neurons in the central complex of insects. Furthermore, we show that this inhibitory network in the aHB forms highly organized reciprocal connections in the dorsal interpeduncular nucleus (dIPN). This architecture agrees with the connectivity scheme required to support ring attractor models of head direction networks (Skaggs et al., 1995; Zhang, 1996) and can provide the substrate for a cognitive map in this vertebrate brain.

## Results

### A population of cells with ring attractor dynamics in the fish aHB

We performed volumetric lightsheet imaging in 7 dpf to 9 dpf zebrafish larvae expressing GCaMP6s in GABAergic neurons in the aHB (Figure 1, Supplementary Figure 1a).

**Figure 1:**
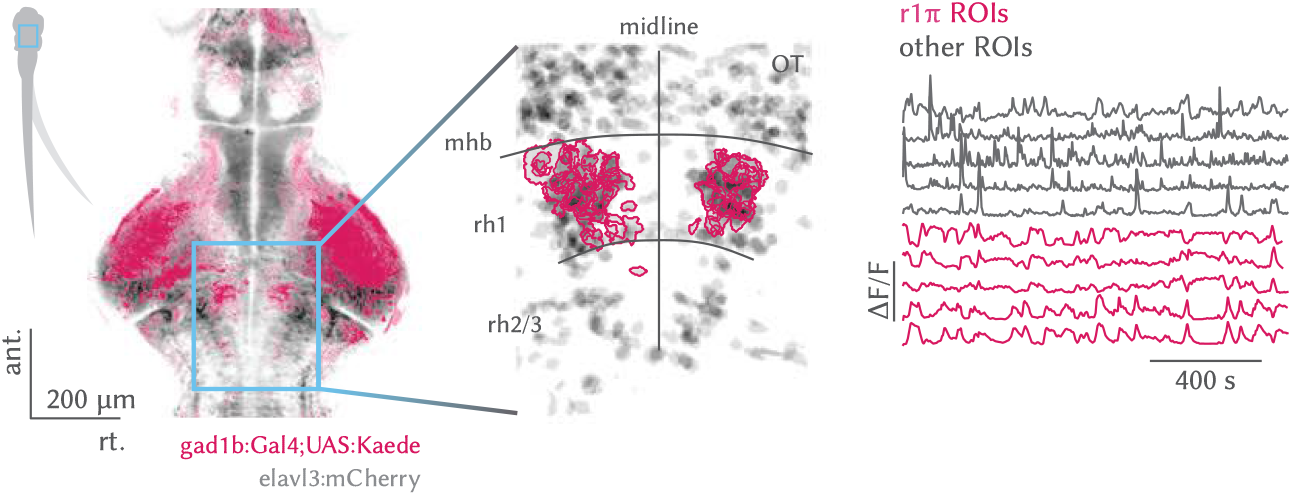
*Left*, expression pattern of the *Tg(gad1b:Gal4)* line over a brain reference. *Center*, example view from an imaging experiment. r1π region of interests (ROIs) are highlighted over the shades of all ROIs from the experiment (OT: *optic tectum*, mhb: *midbrain/hindbrain boundary*, rh: *rhombomere*). *Right*, example traces from one experiment.

Larvae were head-restrained but free to move their tail, and were imaged either in darkness or while presented with a visual stimulus in either closed or open-loop (see *Visual stimuli and experimental groups*).

We observed a population of 50-100 neurons (median = 74, Q1 = 48, Q3 = 115, n = 31 fish) with a sustained bump of activity propagating either clockwise or counterclockwise across the network in a horizontal plane (Figure 2, Movie 1).

**Figure 2:**
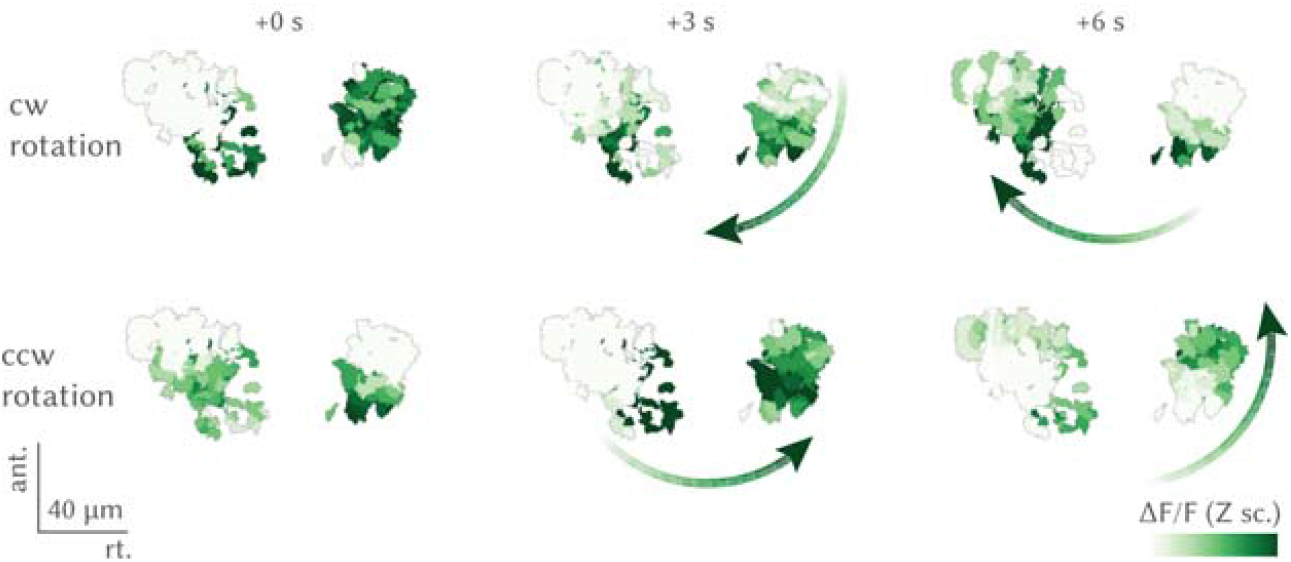
Circular propagation of activity. Intensity of fluorescence for all ROIs in the course of a *top*, clock-wise and *bottom*, counterclockwise propagation event. The arrow shows the direction of activity propagation.

**Movie 1:**
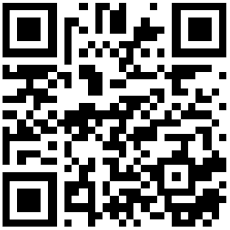
https://doi.org/10.6084/m9.figshare.17871875

These GABAergic neurons were located in rhombomere 1 consistently across fish (Supplementary Figure 1b-c). In order to further characterize the dynamics of the network, we performed principal component analysis (PCA) and observed that the first two principal components (PCs) captured over 80% of the variance (median = 0.800, Q1 = 0.770, Q3 = 0.836, n = 31 fish) (Supplementary Figure 2a-b). Moreover, the trajectory in the phase space defined by the first two PCs was constrained to a circle over the whole duration of the experiment which lasted tens of minutes (Figure 3). For their location in rhombomere 1, the fact that they have an anticorrelated partner at a πangle on the PC space, and based on their morphological features described in a later section of the paper, we call those cells **r1π neurons**.

**Figure 3:**
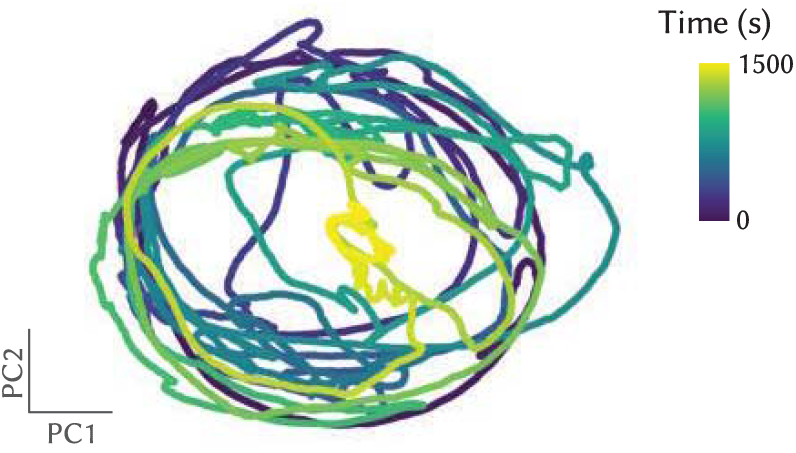
Trajectory in 2d PCs phase space of the network, color-coded by time.

To visualize how the activity of r1π neurons and their anatomical location was related, we projected the activity of each neuron onto a two dimensional subspace, by performing a different PC projection now over the time axis. When projected over the first two principal components (variance explained: median = 0.858, Q1 = 0.827, Q3 = 0.868, n = 31 fish; Supplementary Figure 1c-d), r1π neurons were organized in a circle, with the angle around the circle, *α*, correlating with the neuron’s anatomical location (Fisher-Lee circular correlation *ρ*_*t*_ : median = 0.549, Q1 = 0.298, Q3 = 0.696, n = 31 fish; Figure 4, Supplementary Figure 3). This matches the observation from the raw data of a bump of activity propagating across the network: the circular dynamics we observe in phase space can be seen to correspond to the activity propagation across an anatomical circle of neurons.

**Figure 4:**
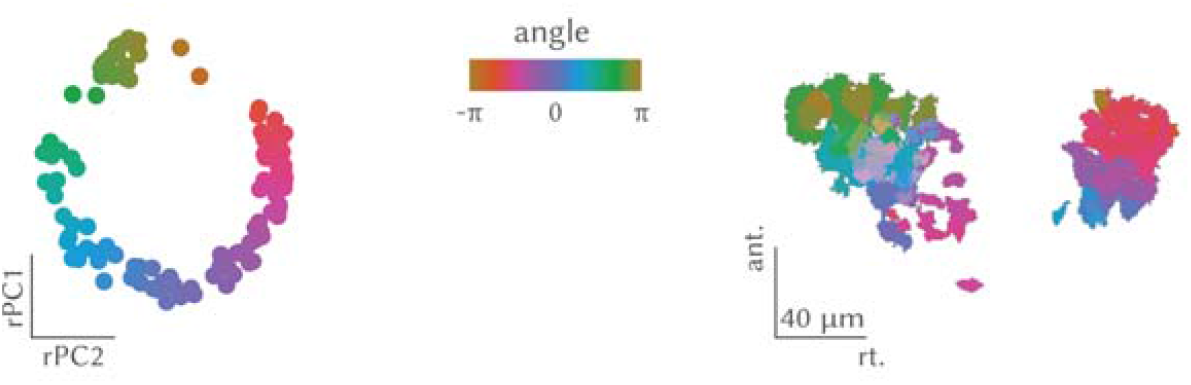
*Left*, Projection over the first 2 rotated principal components (rPCs) in time of all the r1π neurons, color-coded by angle around the circle (for rPC calculation, see *Rotated principal component calculation*, and Figure 27). *Right*, Anatomical distribution of the same neurons, color-coded by angle in rPC space.

In order to describe the position of the bump of activity within the network at any instance in time, we defined an instantaneous network phase *ϕ*(*t*) as the average over neurons of their angle *α* defined above, weighted by their activity at time *t* (Figure 30 and Movie 2). This network phase *ϕ*(*t*) described the angle along the circular trajectory in the network phase space (Supplementary Figure 4a). We anchored *ϕ* by setting it to be 0 when the posterior part of the ring was active, and to increase with clockwise rotations of activity in the horizontal plane (see *Rotated principal component calculation*, Supplementary Figure 3a-c).

To visualize the evolution of the network activity over time, the traces of neurons were sorted by their angle *α* (Figure 5 and Supplementary Figure 5). In this visualization, it can be observed how the phase marks the position of the bump peak as it translates across the network.

**Figure 5:**
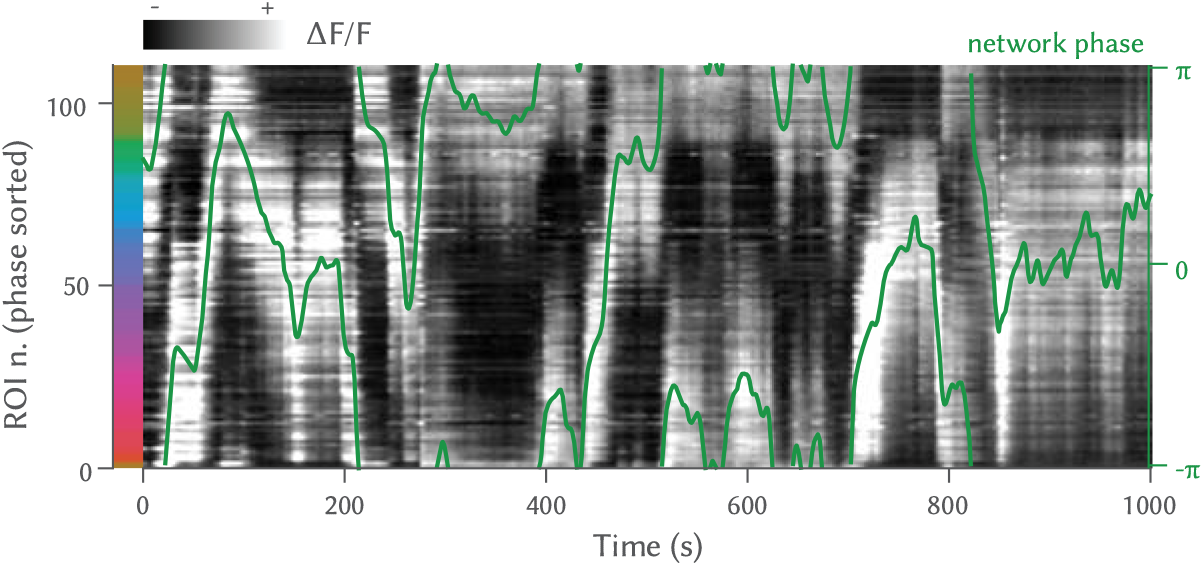
Traces of r1π neurons, sorted by angle in PC space for the neurons, and phase of the network.

To further characterize how individual neurons contributed to the network activity, we computed the average over time of the network activity profile (see *Calculation of average activity profile*, Figure 32). We observed that it was close to a sine wave, with a full width at half maximum (fwhm) of approximately π (mean = 2.910 ± 0.115 rad, n = 31 fish) over the circle of neurons (Supplementary Figure 6a,b). When looking at the tuning curves of individual neurons over network phase, they had an approximately sinusoidal shape, with fwhm of ≈ *π* (Supplementary Figure 4b).

**Movie 2:**
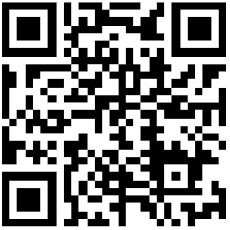
https://doi.org/10.6084/m9.figshare.17871941

### The r1π network integrates heading direction

We next investigated what was driving changes in the phase of the network. We observed that the phase was stable in epochs when the fish was not moving, and was changing the most during sequences of left or right swims (Figure 6).

**Figure 6:**
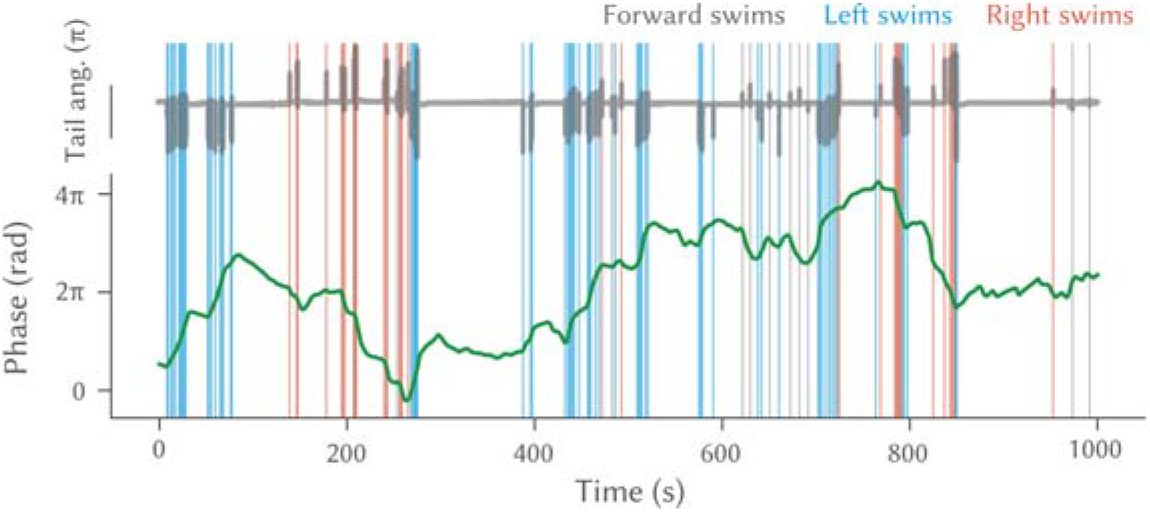
Network phase and motor activity. *Top*, Tail angle over time. Vertical lines mark the occurrence of swims. *Bottom*, Unwrapped network phase over time.

Moreover, sequences of left or right turns were accompanied by clockwise or counterclockwise rotations respectively of the network phase (activity), irrespective of the starting phase position (Figure 7 and Supplementary Figure 7a).

**Figure 7:**
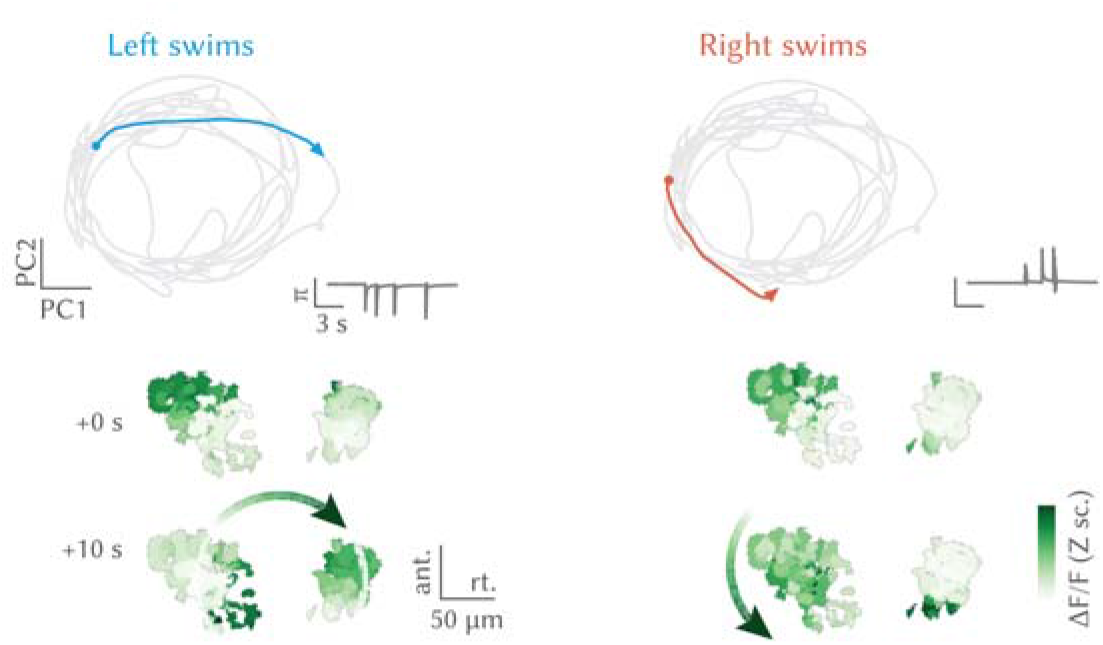
Network trajectory during sequences of left and right swims. *Top, left*, Trajectory in phase space during a sequence of left swims (see tail angle in the insert). *Bottom, left*, State of activation of the network before and after the sequence. *Right*, The same plots for a sequence of right swims.

To quantify this relationship, we computed the swim-triggered change in phase, and we noticed that it was consistently increasing or decreasing after left and right swims (Figure 8), so that left swims (counterclockwise rotations of the fish) would produce clockwise rotations in the network, and right swims (clockwise rotations of the fish) would produce counter-clockwise rotations in the network; forward swims did not produce any consistent change in the network phase.

**Figure 8:**
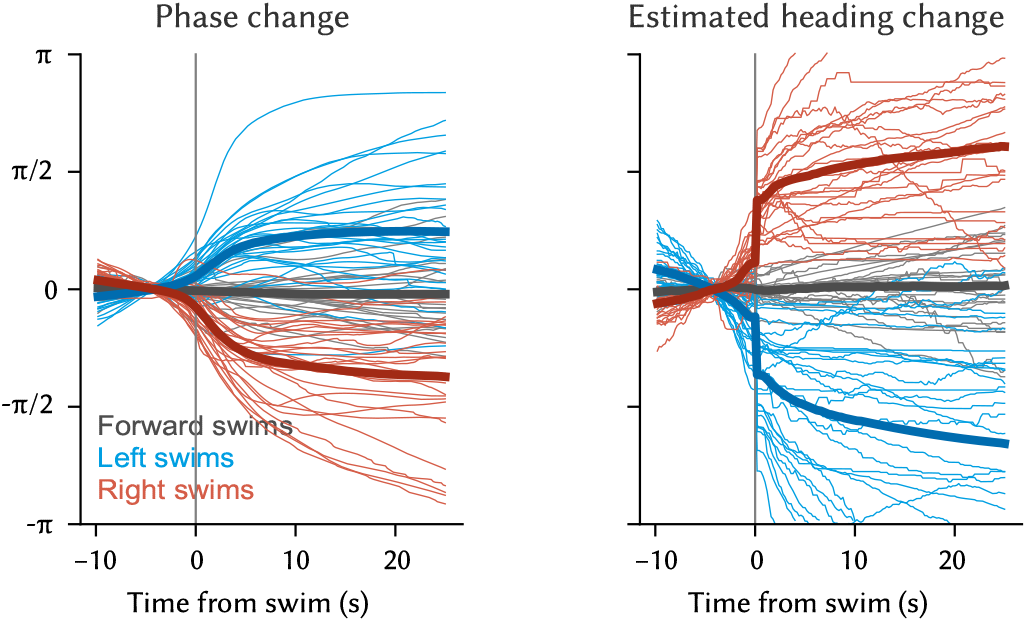
*Left*, Swim-triggered average change in network phase for all fish (thin lines, n = 31 fish) and their average (thick lines). *Right*, Swim-triggered average change in estimated heading direction for all fish (thin lines, n = 31 fish) and their average (thick lines).

Importantly, we observed that the probability for the network phase to be in any state between -π and π when a swim occurred was not different for left, right, and forward swims (Supplementary Figure 7b). This indicates that the absolute/instantaneous network phase does not correlate with specific behavioral outputs.

The swim-triggered changes in network phase (Figure 8, left) show that the amount of angular change elicited by a single swim after 10 s was approximately *π*/4 (median = 0.828 rad, Q1 = 0.492, Q3 = 1.28, n = 31 fish), comparable in size to the angle turned by a swim performed by a freely swimming fish (Figure 8, right) (Huang et al., 2013). Moreover, continuous turning in one direction resulted in several rotations around the network (Supplementary Figure 7c). We therefore hypothesized that the network could work as a heading direction integrator, shifting the position of its activity with every turn and keeping track of the heading direction of the animal in allocentric coordinates as schematized in Figure 9.

**Figure 9:**
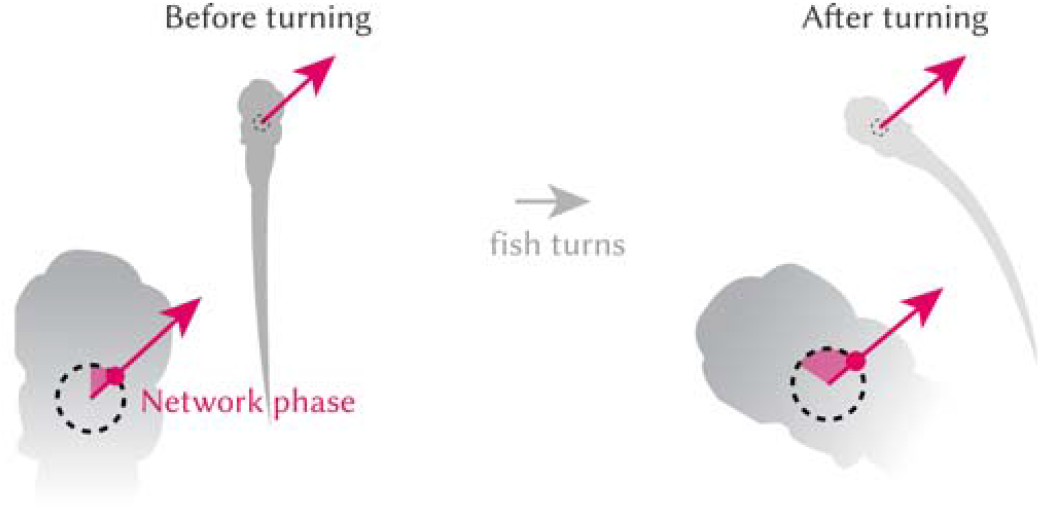
Schematic to show how the network phase changes during a turn.

To understand to which degree the network could produce an estimate of heading direction over time, we reconstructed a fictive heading direction for the head-embedded fish integrating the angle turned by each swim over time (see *Estimated heading calculation and correlation with phase*, Figure 33). This reconstructed heading direction and the network phase were significantly anticorrelated over a period of minutes (Figure 10).

**Figure 10:**
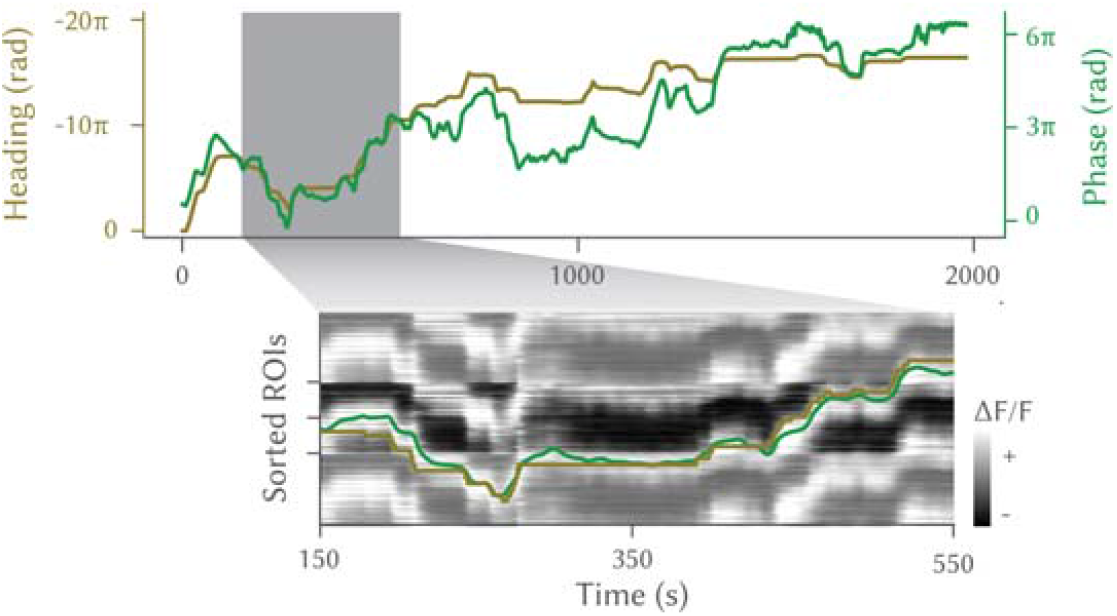
*Top*, Network phase and estimated fish heading for the entire duration of an experiment. Note that the axes are different, and have opposite signs. *Bottom*, The enlargement shows the same traces, overlaid on the traces from the r1π neurons, tiled to match the phase unwrapping.

Although some errors accumulated over time, around each time point the phase in the network could be used to read out an estimate of the current heading direction (Figure 10, enlargement). In fact, the two were significantly anticorrelated (correlation r: median = −0.723, Q1 = −0.863, Q3 = −0.564, n = 31 fish; Figure 11, Supplementary Figure 8).

**Figure 11:**
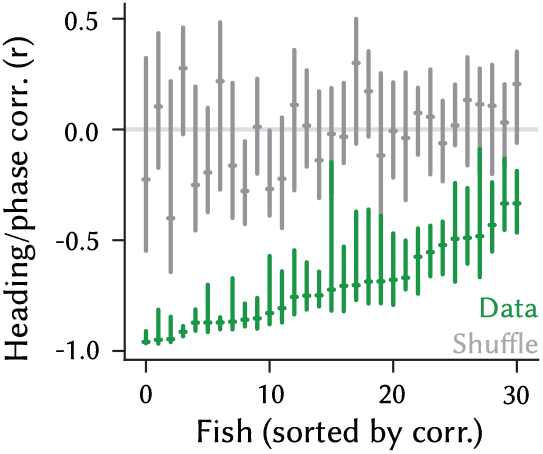
Correlation of heading and network phase for all fish in the dataset, compared with a shuffle of the same data. Bars report median and first and third quartile for the data and a shuffle calculated over 5 min intervals across the experiment (*P* < 0.01 for each of n = 31 fish).

### The r1π network is not affected by visual inputs

Next, we asked whether sensory inputs are required for the observed heading direction integration. As our preparation was head restrained (Figure 12, Figure 22), we could ascertain that vestibular sensory inputs were not required, even though these are known to contribute to the mammalian heading direction system (Yoder and Taube, 2014).

**Figure 12:**
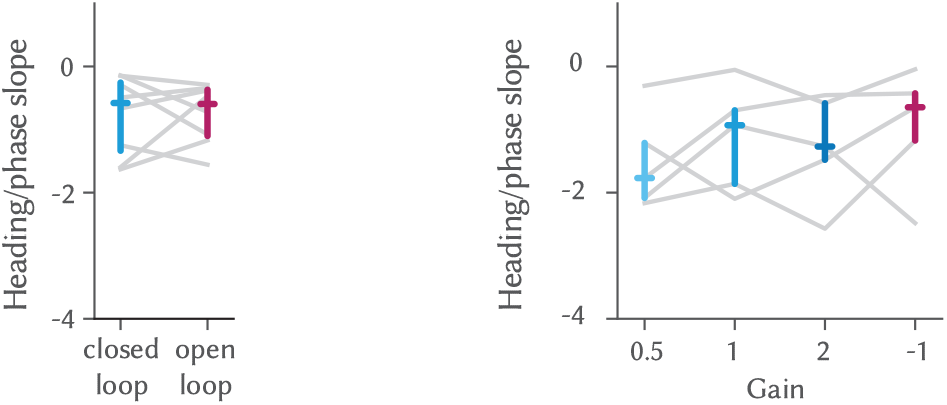
*Left*, Slope of the regression between estimated heading and network phase in closed-and open-loop epochs (not significant difference, Wilcoxon test, n = 8 fish). *Right*, Slope of the regression between heading and network phase in different gain conditions; comparison was not significant between any of the conditions (uncorrected Wilcoxon test, n = 5 fish).

In our experiments, we observed the integration of heading direction in both complete darkness and in open-loop (without visual reafference) (Figure 12, center and Supplementary Figure 9a), indicating that visual feedback is not required for a stable heading direction representation. Furthermore, we tested closed-loop experiments with a range of gains that provided different amounts of visual feedback. In these experiments, we observed no relationship between the representation of heading direction and the experimental gain (Figure 12, right and Supplementary Figure 9b). This shows that visual feedback not only is not required, but that it does not contribute to the activity we observe, and suggests that efference copies are its main driver.

Interestingly, the activity of left and right GABAergic clusters in rhombomeres 2 and 3, immediately caudal to the r1π neurons, show a remarkable degree of correlation with leftward and rightward swims, respectively (Figures 13 and 35 and Supplementary Figure 10). These neurons might provide the motor efference input to the r1π network.

**Figure 13:**
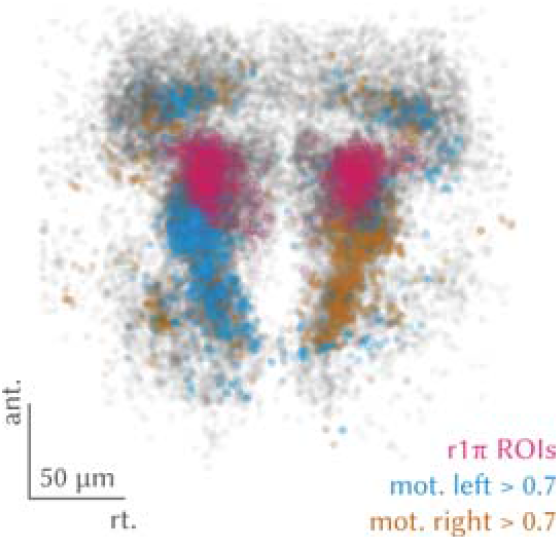
Distribution of directional swim-related ROIs.

### The r1π network is modulated by eye movements

Subpopulations of cells in the aHB are known to represent eye-related variables such as eye position and saccade timing (Wolf et al., 2017; Ramirez and Aksay, 2021). We therefore decided to understand whether eye motion could also modulate the phase of the network. To this end, we freed the eyes of the larvae in a subset of experiments and tracked their motion together with that of the tail. In periods where swimming was absent, we observed that eye motion could explain some low amplitude modulation in the network phase (Figure 14 and Supplementary Figure 11d), although eye motion on its own did poorly compared to heading direction when swimming did occur (Supplementary Figure 11a). Interestingly, the sign of the modulation was consistent with the heading changes, with leftward saccades increasing the network phase as leftward swims do, and rightward saccades decreasing it as rightward swims do.

**Figure 14:**
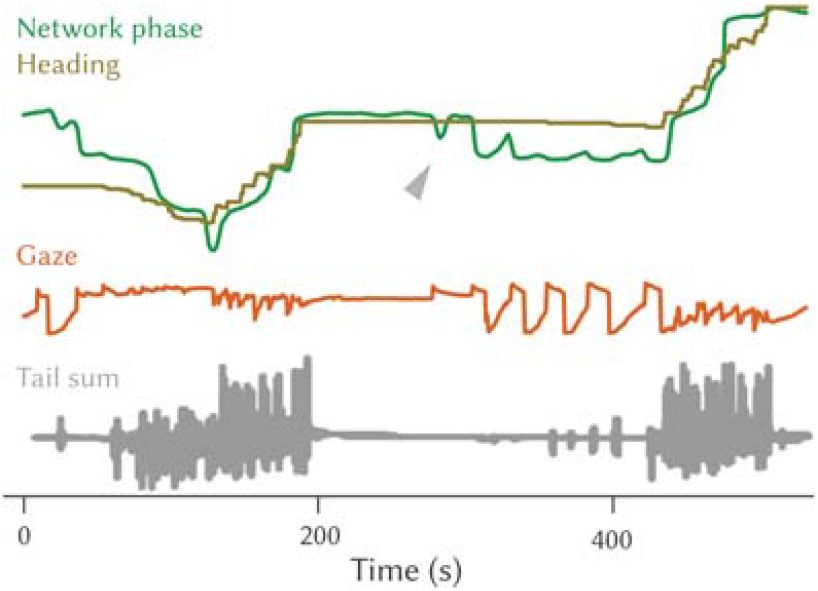
Network phase represented with (flipped) heading direction and gaze direction. The arrow highlights a period of very sparse swimming and large saccades.

### aHB neurons arborize in the dorsal IPN

We proceeded to investigate the anatomy of neurons in the aHB. Anatomical stacks of the GABAergic line we used show a prominent, bilaterally-paired, tract of fibers that extended ventro-medially from the GABAergic nuclei of rhombomere 1 towards the dorsal IPN (dIPN) (Figure 15, red arrow, and Supplementary Figure 12a,b).

**Figure 15:**
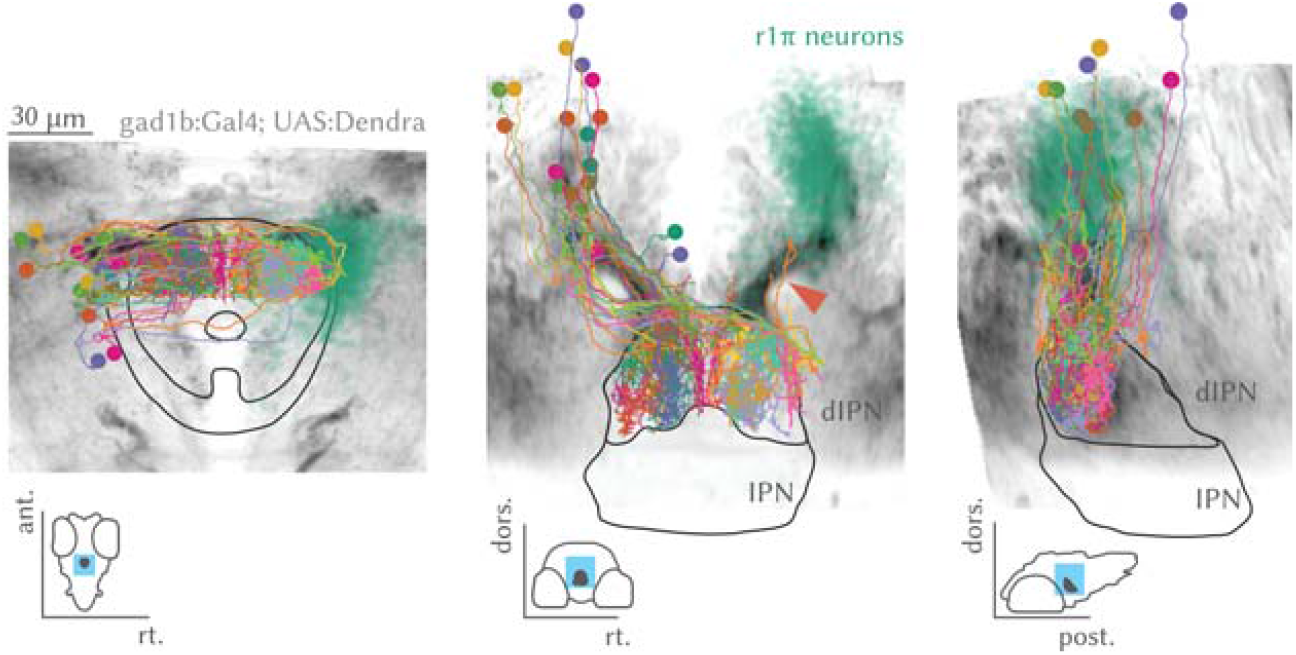
Anatomical projections of a stack from the *Tg(gad1b:Gal4)* line used in the experiments. The lines mark the IPN and dIPN boundaries, and the insets show the position in the brain of the IPN mask and the views. The arrow highlights the tract of fibers that extend from the aHB to the IPN. The r1π neurons from the imaging experiments are shown in the same coordinate space in green on the right, together with the morphology of all neurons reconstructed from the SBEM data on the left.

**Figure 16:**
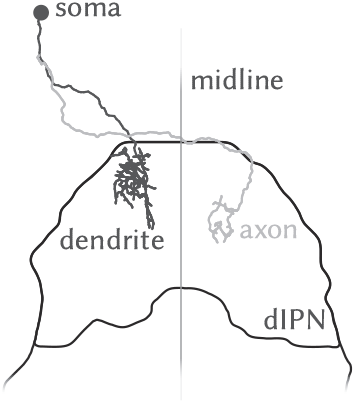
One of the neurons from Figure 15, singled out to show the cell morphology with a process splitting in an ipsilateral dendrite and a contralateral axon.

To reconstruct individual neurons at high resolution, we traced neurons and their projections in a serial block-face electron microscopy (SBEM) dataset. We identified a class of neurons with the somata in the aHB that extended a single projection that bifurcated into a dendrite and axon which ended in the dIPN (Figure 15, Movie 3 and Supplementary Figure 12c).

The small dendritic tree covered a localized compartment in the ipsilateral IPN, whereas the axon projected contralaterally with minimal branching that occurred only in the terminal sections (Supplementary Figure 13).

**Movie 3:**
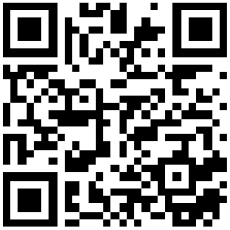
https://doi.org/10.6084/m9.figshare.19608204

To confirm that r1π neurons project to the dIPN, we imaged the same GABAergic line under a two-photon microscope to investigate neuropil activity. Performing the same analysis as for the r1π neurons presented in Figures 4, 5 and 10 uncovered a set of ROIs that were mostly restricted to the dIPN, showed stable circular dynamics and displayed the same relationship to heading direction (Figure 17 and Supplementary Figure 14).

**Figure 17:**
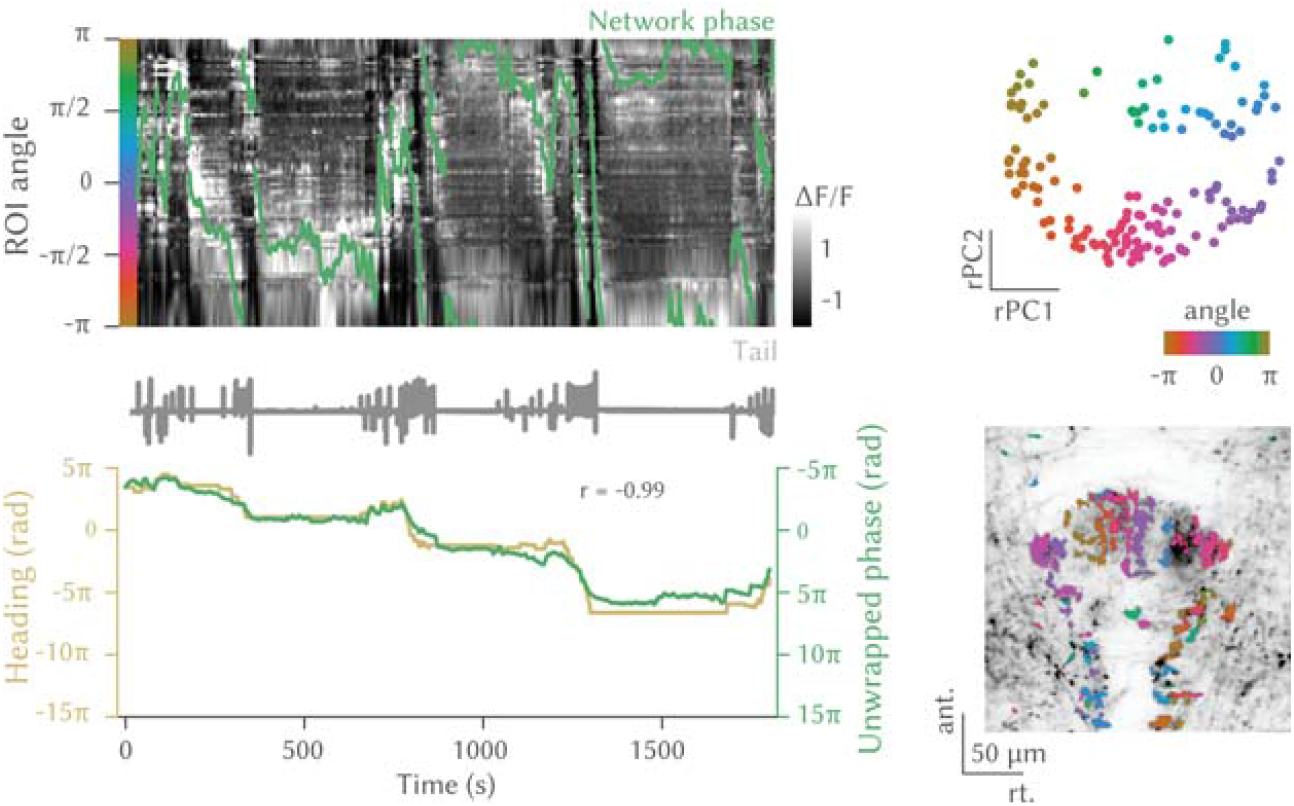
*Left, top*, Traces of ROIs in the dIPN showing r1π-like dynamics, sorted by angle in PC space, and phase of the network (green line). The tail trace is shown in gray on top. *Left, bottom*, Estimated heading direction and network phase are highly correlated. *Right, top*, projection over the first 2 PCs in time of all the ROIs showing r1π- like activity, color-coded by angle around the circle. *Right, bottom*, Anatomical distribution of the same neurons, color-coded by angle in PC space. The anatomy of the recorded plane is shown in the background.

### aHB projections map linearly the functional topology of the ring in the dIPN

It has been suggested that heading directions systems are neuronal implementations of ring-attractor networks, where excitatory activity between neighboring cells is stabilized and localized by long-range inhibitory connections. We therefore wanted to investigate whether there is any evidence that the morphology and projections of the GABAergic r1π neurons could implement such a structure. To this end, we turned back to the SBEM reconstructions. We observed that the projections of different neurons occupy different locations in the medio-lateral axis and appear to cover the whole dIPN (Figure 18, Movie 3 and Supplementary Figure 13). Moreover, the distance of the dendrite from the midline anticorrelated with the distance of the axon from the midline (r = −0.9,n = 19 neurons), meaning that a neuron with a lateral dendrite would extend a medial axon and vice-versa (Figure 18 *left*, Supplementary Figure 15a). In addition, given the anatomical organization of the activity we observed in the aHB, we would expect a correlation between the antero-posterior position of a cell’s soma and the distance of its dendrites from the midline. This is in fact what we observed (r = −0.65, n = 19 neurons) (Figure 18 *right*, Supplementary Figure 15).

**Figure 18:**
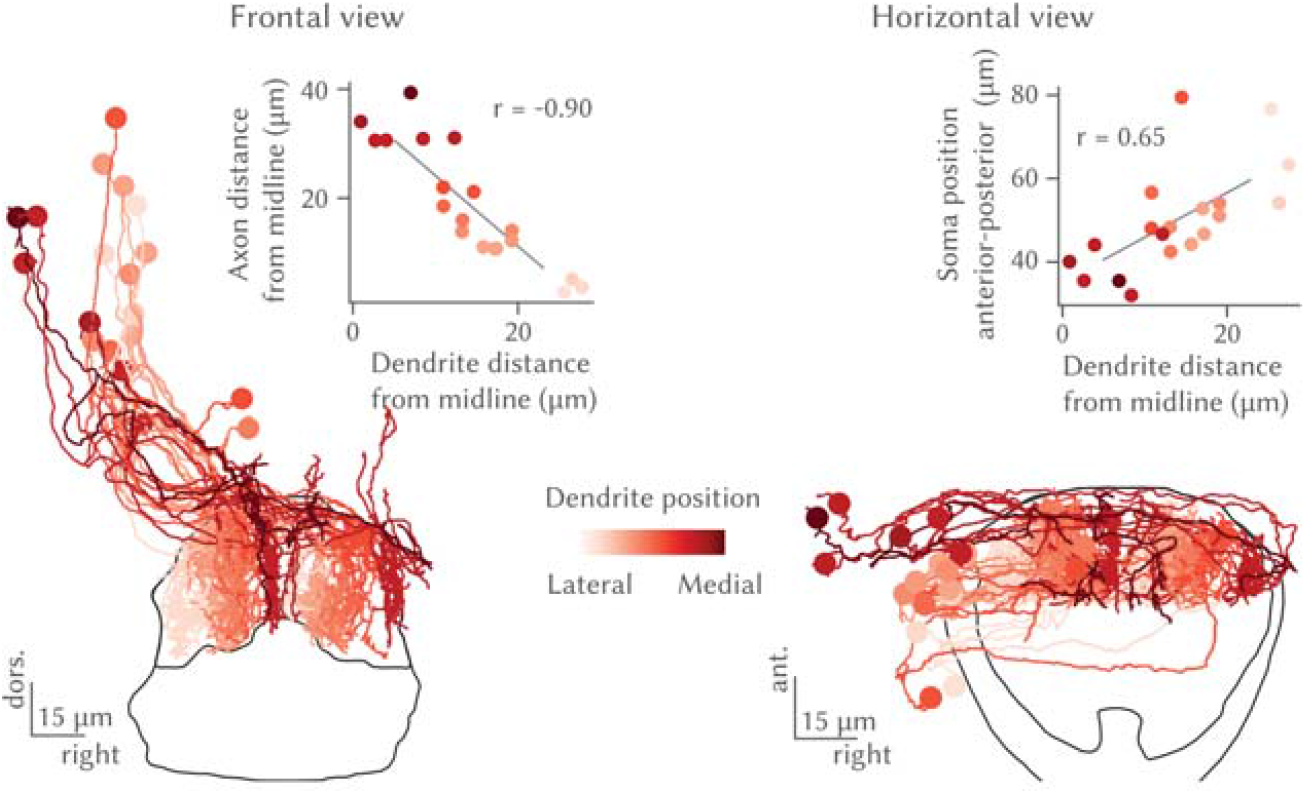
*Left*, Frontal view of IPN-projecting aHB neurons, color-coded by position of the dendrite on the coronal axis (lateral-medial). Inset: scatterplot of the distance from the midline of dendrite and axon for each neuron (R = −0.90, n = n = 19 neurons). *Right*, Horizontal view IPN-projecting aHB neurons, color-coded by position of the dendrite on the coronal axis (lateral-medial). Inset: scatterplot of the distance from the midline of dendrite and position of the soma of the anteroposterior axis for each neuron (r = −0.65, n= 19 neurons).

Such an organization would predict that in the activity recorded from the dIPN, pixels that are the most correlated with each other are at a fixed distance on the medio-lateral axis, as their signal comes from the dendrites and axons of the same neurons. Indeed, we found this type of pattern when examining data from single fish (Figure 19) and across all fish (Figure 20 *top*, Supplementary Figure 16). The distance of the side lobes observed in the functional correlation correlated with the distance of each neuron’s axon from its dendrite (Figure 20, bottom, Supplementary Figure 15c). These observations suggest that a circular functional structure in the aHB, corresponding to angles from −*π* to *π*, is coupled to a linear structure in the dIPN (Figure 21, *left*). Neurons whose soma are at opposite sides of the circular organization of aHB target with their axons their respective dendrites (Figure 21, *right*). In this way, a neuron is ideally placed to inhibit its corresponding out-of-phase neurons. This projection pattern could stabilize and localize the heading direction activity we observed in the aHB network by providing long range inhibition.

**Figure 19:**
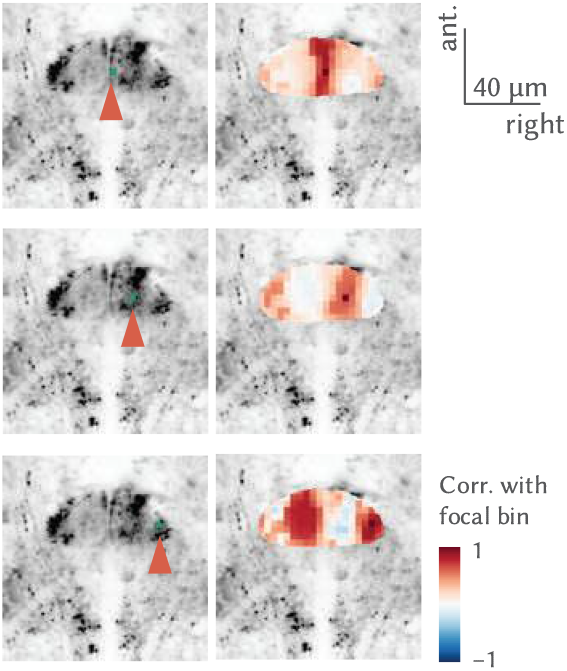
*Left*, Anatomy of *Tg(gad1b:Gal4, UAS:GCaMP6s)* from a two photon experiment with a focal bin highlighted by the red arrow and *right*, maps of correlations of bins with the focal bin. Each row corresponds to a different focal point.

**Figure 20:**
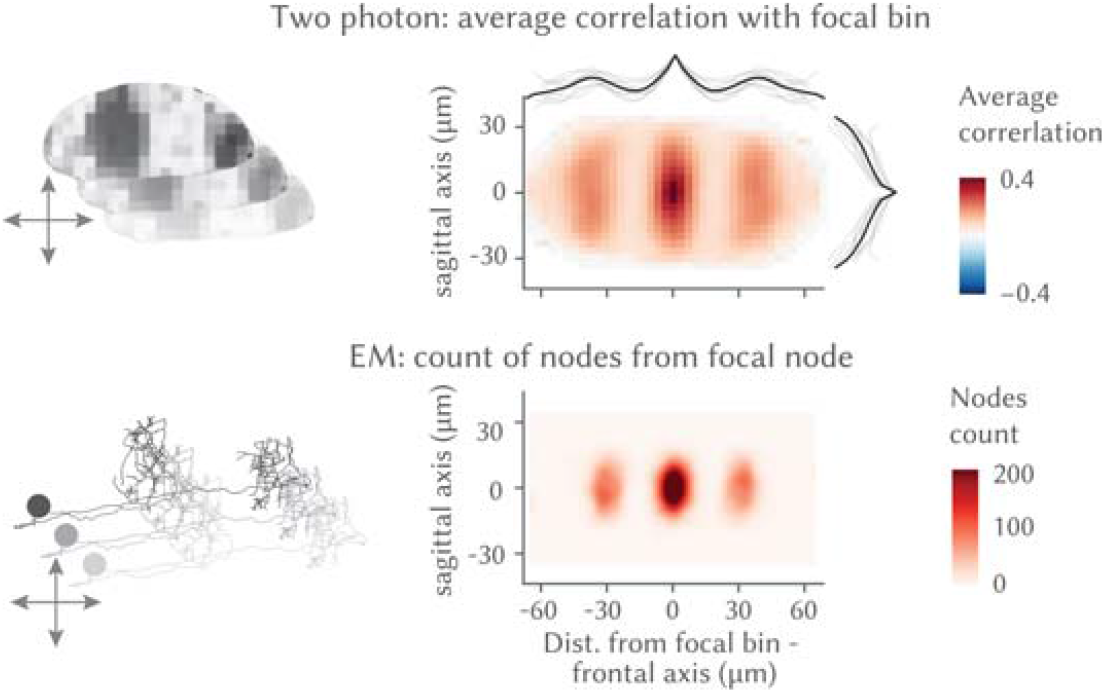
*Top*, Average correlation of bins at different distances around a focal bin in the two-photon data. The lines on the side show individual fish means across each axis (thin lines) and population average (thick line). *Bottom*, Count of nodes around a focal node in the SBEM data.

**Figure 21:**
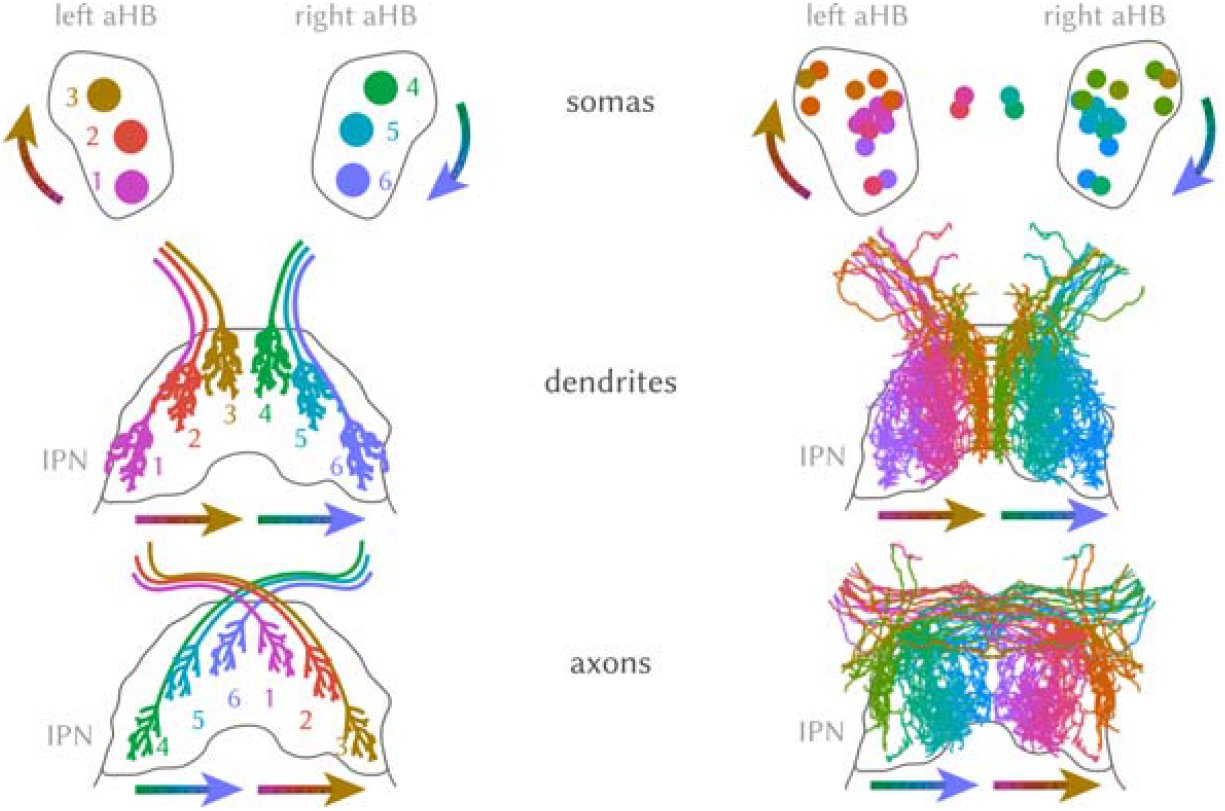
*Left*, Schematics of the organization of aHB neuron somata (top) and their dendritic (middle) and axonal (bottom) projections in the IPN. *Right*, The same but with actual reconstructed EM neurons that have been mirrored on both sides. The color code is based on the position of the dendrite.

**Figure 22:**
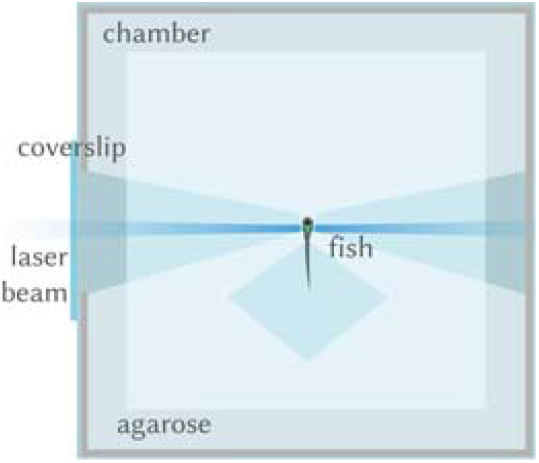
Schema of the preparation for the lightsheet, with a fish head-restrained in agarose in the chamber.

**Figure 23:**
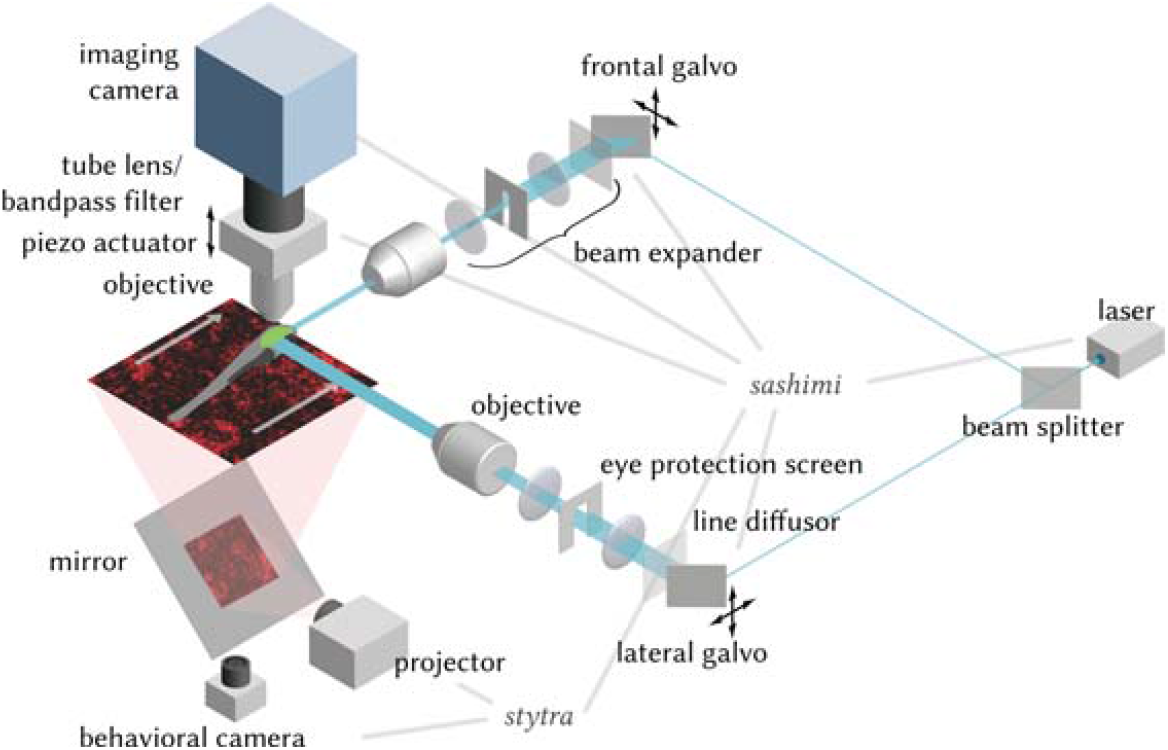
Schema of the lightsheet microscope described in the text. Software is indicated in italics, and gray lines indicate the parts of the setup controlled by each program.

## Discussion

In this study, we describe a network of GABAergic neurons in rhombomere 1 of the larval zebrafish hindbrain that encodes the heading direction of the fish in external (allocentric) coordinates. As this representation persists even in the absence of external landmarks and salient sensory stimuli, it is likely generated by the integration of efference copies. We show that motor-related activity exists nearby that may be serving this purpose, in agreement with previous reports (Dunn et al., 2016; Chen et al., 2018). This observation confirms the existence of an internal model of turning in the zebrafish brain.

Remarkably, the heading direction is represented by a bump of activity that propagates clockwise and counter-clockwise in the horizontal plane with leftward and rightward movements of the fish. This is the first evidence of an anatomical organization for the heading direction system in a vertebrate, and suggests the existence of simple topographical principles in the wiring of the network. Indeed, our electron microscopy reconstructions suggest that heading direction neurons of the aHB connect to each other in a precise way, with neurons whose functional activation is in complete antiphase making reciprocal connections in the dIPN. Based on their location, on their consistent antiphase projection pattern, we term those cells r1π heading direction neurons.

In mammals, the GABAergic (Allen and Hopkins, 1989; Wirtshafter and Stratford, 1993) dorsal tegmental nucleus is considered to be one of the earliest subcortical structures within the heading direction pathway, and tracing studies have identified reciprocal connections between the DTN and the IPN (Contestabile and Flumerfelt, 1981; Groenewegen et al., 1986; Liu et al., 1984). The heading direction neurons that have been observed in the DTN are broadly tuned (Sharp et al., 2001), similarly to the neurons we report in the fish aHB. Moreover, tegmental afferents to the mammalian (rostral) IPN form highly compartmentalized arborizations (Herrick, 1948; Iwahori et al., 1993), like we observe in the reconstructions of aHB fibers.

Theoretical studies (Skaggs et al., 1995; Zhang, 1996; Hansel and Sompolinsky, 1998; Hulse and Jayaraman, 2020) proposed the notion of ring attractor networks as a mechanism to encode heading direction information, and evidence of ring attractor-like dynamics has been found in the rodent heading direction system (Chaudhuri et al., 2019; Clark et al., 2009). However, a mechanistic understanding based on the neuronal connectivity that underlies such dynamics is still missing in vertebrates.

This link between function and structure exists in the insect central complex, where elegant studies have described networks that encode heading direction and constitute a neuronal implementation of a ring attractor network (Seelig and Jayaraman, 2015; Green et al., 2017; Kim et al., 2017; Fisher et al., 2019; Suver et al., 2019; Lyu et al., 2022). The level of detail being uncovered in these circuits allows for a mechanistic understanding of how a brain can integrate external and internal sensory cues, efference copies and carry out coordinate transformations that are important for behavior. The network we observe bears intriguing similarities with this system and a detailed comparative analysis of both may uncover important theoretical insights into persistent neuronal representations in general and head-direction systems and ring attractors in particular (Hulse and Jayaraman, 2020).

We still need to address how external cues influence this network activity to determine whether the “North” of the representation points in a behaviorally relevant direction in space. Previous work has shown that zebrafish larvae might use an internal representation of heading direction to efficiently reorient in a phototaxis task (Chen and Engert, 2014), and the interpeduncular nucleus can be implicated in zebrafish directional behavior (Dragomir et al., 2020; Cherng et al., 2020). Information from external cues, as well as strong excitatory drive could be provided by the dense projections from excitatory habenular neurons (Hong et al., 2013), which could form synapses with an all-to-all connectivity with the dendrites of the heading direction neurons (Bianco and Wilson, 2009). Although previously overlooked, the aHB-IPN circuit could provide an inroad to understanding the mechanisms underpinning cognitive maps in vertebrates.

## Materials and Methods

### Zebrafish husbandry

All procedures related to animal handling were conducted following protocols approved by the Technische Universität München and the Regierung von Oberbayern. Adult zebrafish (*Danio rerio*) from Tüpfel long fin (TL) strain were kept at 27.5 °C to 28 °C on a 14 /10 light cycle, and hosted in a fish facility that provided full recirculation of water with carbon-, bio- and UV filtering and a daily exchange of 12% of water. Water pH was kept at 7.0 to 7.5 and conductivity at 750 μSv to 800 μSv. Fish were hosted in 3.5 l tanks in groups of 7 to 10 animals. Adults were fed with Gemma micron 300 (Skretting) and live food (*Artemia salina*) twice per day and the larvae were fed with Sera micron Nature (Sera) and ST-1 (Aquaschwarz) three times a day.

All experiments were conducted on 6 dpf to 9 dpf larvae of yet undetermined sex. The week before the experiment, one male and one female or three male and three female animals were left breeding overnight in a Sloping Breeding Tank or breeding tank (Tecniplast). The day after, eggs were collected in the morning, rinsed with water from the facility water system, and then kept in groups of 20 to 40 in 90 cm Petri dishes filled with 0.3× Danieau’s solution (17.4 mM NaCl, 0.21 mM KCl, 0.12 mM MgSO4, 0.18 mM Ca(NO3)2, 1.5 mM hepes, reagents from Sigma-Aldrich) until hatching and in water from the fish facility afterwards. Larvae were kept in an incubator that maintained temperature at 28.5 °C and a 14 /10 light/dark cycle, and their solution was changed daily. At 4 dpf to 5 dpf, animals were lightly anesthetized with Tricaine mesylate (Sigma-Aldrich) and screened for fluorescence under an epifluorescence microscope. Animals positive for GCaMP6s, Dendra or mCherry fluorescence were selected for the imaging experiments.

### Transgenic animals

The *Tg(gad1b/GAD67:Gal4-VP16)mpn155* (referred to as *Tg(gad1b:Gal4)*) was used for all experiments, which drives expression in a subpopulation of GABAergic cells under *gad1b* regulatory elements (Förster et al., 2017). The animals for functional imaging and anatomical experiments were double transgenic with *Tg(UAS:GCaMP6s)mpn101* (Thiele et al., 2014) and *Tg(UAS:Dendra-kras)s1998t* (Arrenberg et al., 2009), respectively. In some anatomical experiments, the animals also had *Tg(elavl3:H2B-mCherry)*, which was generated by Tol2 transposon-mediated transgenesis. All the transgenic animals were also *mitfa-/-* and thus lacked melanophores (Lister et al., 1999).

### Lightsheet experiments

#### Preparation

For lightsheet experiments, animals were embedded in 2.2% low-melting point agarose (Thermofisher) in a custom lightsheet chamber. The chamber consisted of a 3D printed frame (`.stl` file link: https://github.com/portugueslab/hardware/blob/master/chambers/lightsheet_chamber_v3.stl) with a glass coverslip sealed on the side in the position where the lateral beam of the lightsheet enters the chamber, and a square of transparent acrylic on the bottom, for behavioral tracking (see *Lightsheet microscope*).

The chamber was filled with water from the fish facility system and agarose was removed along the optic path of the lateral laser beam (to prevent scattering), and around the tail of the animal, to enable movements of the tail (Figure 22). In some larvae, the eyes were also freed from the agarose. After embedding, fish were left recovering 1h to 6h before the imaging session. Before starting the imaging, light tapping on the side of the chamber was used to select the most active fish for the experiment.

#### Lightsheet microscope

Imaging experiments were performed using a custom-built lightsheet microscope. A 473 nm wavelength laser source (modulated laser diodes, Cobolt) was used to produce a ∼1.5 mm laser beam that was conveyed on the excitation scanning arm. The arm consisted in a pair of galvanometric mirrors that scanned vertically and horizontally; a line diffuser (Edmund Optics) to minimize stripe artifacts (Taylor et al., 2018), a 2× telescope composed by a 75 mm and a 150 mm focal distance lens (Thorlabs) that expanded the beam before it entered a low numerical aperture air objective (Olympus) that focused it through the lateral glass coverslip of the lightsheet chamber on the fish. The excitation light sheet was generated by scanning at 800 Hz the beam on the horizontal plane. A paper screen was positioned in the image conjugate plane within the telescope lens pair to protect the eyes of the fish from the lateral scanning of the laser beam. The emitted fluorescence was collected with a 20× water-immersion objective (Olympus), filtered with a 525/50 band-pass filter (AHF Analysentechnik) and focused on a cmos camera (Hamamatsu Photonics) with a tube lens (Thorlabs).

The imaging acquisition was run using `sashimi`, a custom Python-based software (Štih et al., 2022a) to coordinate the laser scanning, the camera triggering and the piezo movement. The objective was moved with a sawtooth profile with a frequency of 5 Hz in most experiments (frequency was adjusted to 3 Hz in experiments where a larger vertical span was scanned).

TTL pulses locked with the scanning profile of the piezo were sent to the camera to trigger the acquisition of each plane at a fixed vertical position during the scanning. No pulse was sent during the descending phase of the scanning, when the objective would cover a large vertical span in a short time. In most experiments a total of 8 planes were acquired over a range of approximately 80 μm to 100 μm, slightly adjusted for every fish. The resulting imaging data had a voxel size of ∼10 μm× 0.6 μm× 0.6 μm, and a temporal resolution of 3 Hz to 5 Hz.

#### Tail/eyes tracking and stimulus presentation

To monitor tail movements during the imaging session, an infrared LED source (RS Components) was used to illuminate the larvae from above. A camera (Ximea) with a macro objective (Navitar) was aimed at the animal through the transparent bottom of the lightsheet chamber with the help of a mirror placed at 45° below the imaging stage. A longpass filter (Thorlabs) was placed in front of the camera. A projector (Optoma) with a red long-pass filter (Kodak Wratten No.25) was used to display visual stimuli; light from the projector was conveyed to the stage through a cold mirror that reflected the projected image on the 45° mirror placed below the stage. The stimuli were projected on a white paper screen positioned below the fish, with a triangular hole that kept the fish visible from the camera. The behavior tracking part of the rig was very similar to the setup for restrained fish tracking described in (Štih et al., 2019).

Frames from the behavioral camera were acquired at 400 Hz and tail movements were tracked online using `Stytra` (Štih et al., 2019) with Stytra’s default algorithm to fit to the tail 9 segments. The *tail angle* quantity used for controlling the closed-loop was computed online during the experiment in the Stytra application as the difference between the average angle of the first two and last two segments of the tail and saved with the rest of the log from Stytra. For eye tracking, a video of the entire acquisition was saved to be analyzed offline (see below).

The stimulus presentation and the behavior tracking were synchronized with the imaging acquisition with a ZMQ-based trigger signal supported natively by `Stytra`.

### Two photon experiments

For two photon experiments, animals were embedded in 2% low-melting point agarose (Thermofisher) in 30 mm petri dishes. The agarose around the tail, caudal to the pectoral fins, was cut away with a fine scalpel to allow for tail movement. The dish was placed onto an acrylic support with a light-diffusing screen and imaged on a custom-built two-photon microscope previously described in (Dragomir et al., 2020) (Figure 24). The custom Python package `brunoise` was used to control the microscope hardware (Štih et al., 2020).

**Figure 24:**
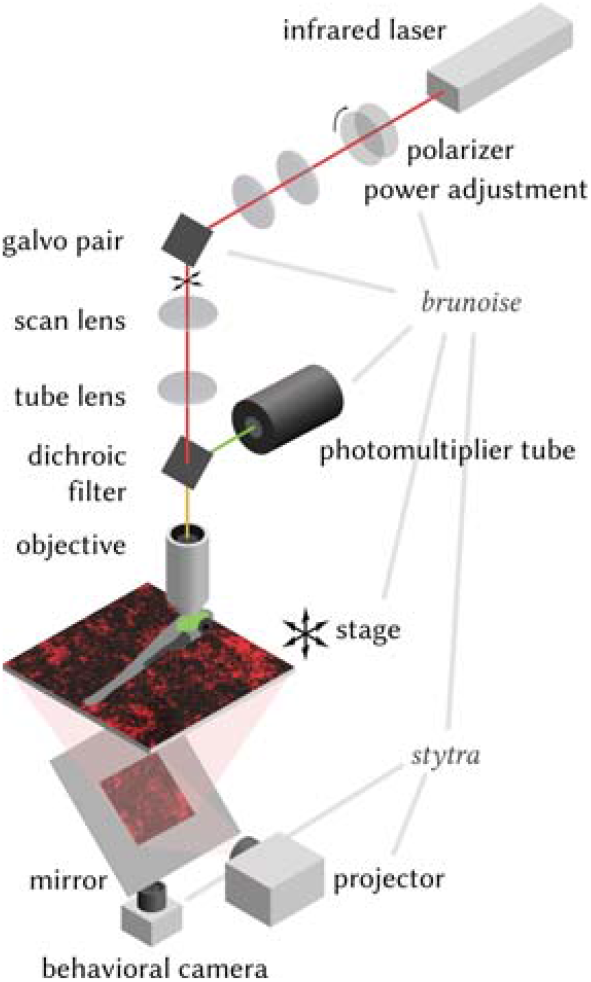
Schema of the two photon microscope described in the text. Software is indicated in italics, and gray lines indicate the parts of the setup controlled by each program.

Full frames were acquired every 334.51 ms in four, 0.83 μm-spaced interlaced scans, which resulted in x and y pixel dimensions of 0.3 μm to 0.6 μm (varying resolutions here depending on field of view covered). After acquisition from one plane was done, the objective was moved downward by 0.5 μm to 4 μm and the process was repeated.

#### Two photon functional experiments

Visual stimuli (see below) were generated using a custom written Python script with the `Stytra` package, and were projected at 60 Hz using an Asus p2e microprojector and a red long-pass filter (Kodak Wratten No.25) to allow for simultaneous imaging and visual stimulation. Fish were illuminated using infrared light-emitting diodes (850 nm wavelength) and imaged from below at up to 200 Hz using an infrared-sensitive charge-coupled device camera (Pike f032pb, Allied Vision Technologies). Tail movements were tracked online using Stytra as described for the lightsheet experiments.

#### Two photon anatomical experiments

High resolution 0.5 μm× 0.5 μm × 0.5 μm two-photon stacks of the aHB and IPN were acquired from fish expressing *gad1b:Gal4* and *UAS:Dendra-kras* transgenes (n = 7 fish, 6 dpf to 7 dpf). The stacks were registered to one another using the Computational Morphometry Toolkit (`CMTK`) (Rohlfing and Maurer, 2003). The transformed stacks were then averaged to generate an average brain stack showing the projections of GABAergic aHB neurons to the IPN.

### Confocal experiments

For confocal experiments, larvae were embedded in 1.5% agarose and anesthetized with Tricaine. Whole brain stacks of three 7 dpf fish expressing *gad1b:Gal4, UAS:Dendra-kras* and *elavl3:H2B-mCherry* transgenes were acquired using a 20× water immersion objective (NA = 1.0) with a voxel resolution of 1 μm× 0.6 μm× 0.6 μm (LSM 880, Carl Zeiss). The stacks were registered to one another using `CMTK` (Rohlfing and Maurer, 2003). The transformed stacks were then averaged to generate an average brain stack showing the expression pattern of the *gad1b:Gal4* on top of pan-neuronal *H2B-mCherry* expression.

### Electron microscopy experiments

#### serial block-face electron microscopy dataset acquisition

Details of the SBEM dataset acquisition will be published elsewhere (Svara et al., *in preparation*). Briefly, a 5 dpf larval *Tg(elavl3:GCaMP5G)a4598* transgenic zebrafish was fixed with extracellular space preservation and stained as described previously (Svara et al., 2018; Briggman et al., 2011). The sample was embedded in an epoxy mixture containing 2.5% Carbon Black (Nguyen et al., 2016). The brain was imaged at a resolution of 14 nm × 14 nm and sections were cut at a thickness of 25 nm. The long axis of each image tile was scanned by gradually moving the stage, while the short axis was scanned with the electron beam. The shape of the tile pattern was determined based on a 4 μm voxel size X-Ray microCT scan (SCANCO Medical AG, Brütisellen) of the embedded sample.

### Visual stimuli and experimental groups

The observations reported in the paper were performed in experiments where different visual stimuli were presented to the fish:

- Darkness: in those experiments, no visual stimuli were presented, the projector was on and a static black frame was displayed;
- Open loop: in open loop epochs, a pink noise pattern was projected and moved in x and theta with a path that was computed from the trajectory of a freely swimming fish taken from a previous experiment in the lab. The stimulus moved backward according to velocity of the fish, and rotated according to changes in its direction. As a result, the fish was presented with the optic flow that it would have perceived moving over a static pink noise pattern with that trajectory.
- Closed loop: a pink noise pattern was projected below the fish; the pattern was static if the animal was not moved, and it translated backward and rotated when the fish performed spontaneous movements. The stimulus moved backward according to an estimate of the velocity of the fish computed using vigor, and rotated according to changes in its direction estimated using the swim bias, so that right turns, i.e, clockwise rotations of the fish, would be matched with clockwise rotations of the stimulus. The gain factor that transformed a given swim bias into an angular velocity was modulated with factors 0.5, 1 and 2 to observe if the slope of the aHB network and the estimated heading would be altered by visual feedback. An additional control gain of −1, where fish would receive a visual feedback opposite to the performed movements, was also included.
- Directional motion: in some experiments, the animal was also shown a pink noise pattern moving in 8 equally spaced directions on the plane, presented one after the other first in clockwise sequence (starting from forward) and then in counter-clockwise sequence.

Visual stimuli were tried in different combinations over different animals in the dataset (see Figure 25). As the activity we describe was not modulated by the presented visual stimuli in those experiments, we pooled together observations from different experimental conditions for all the analyses that quantified the property of the r1π neurons network.

**Figure 25:**
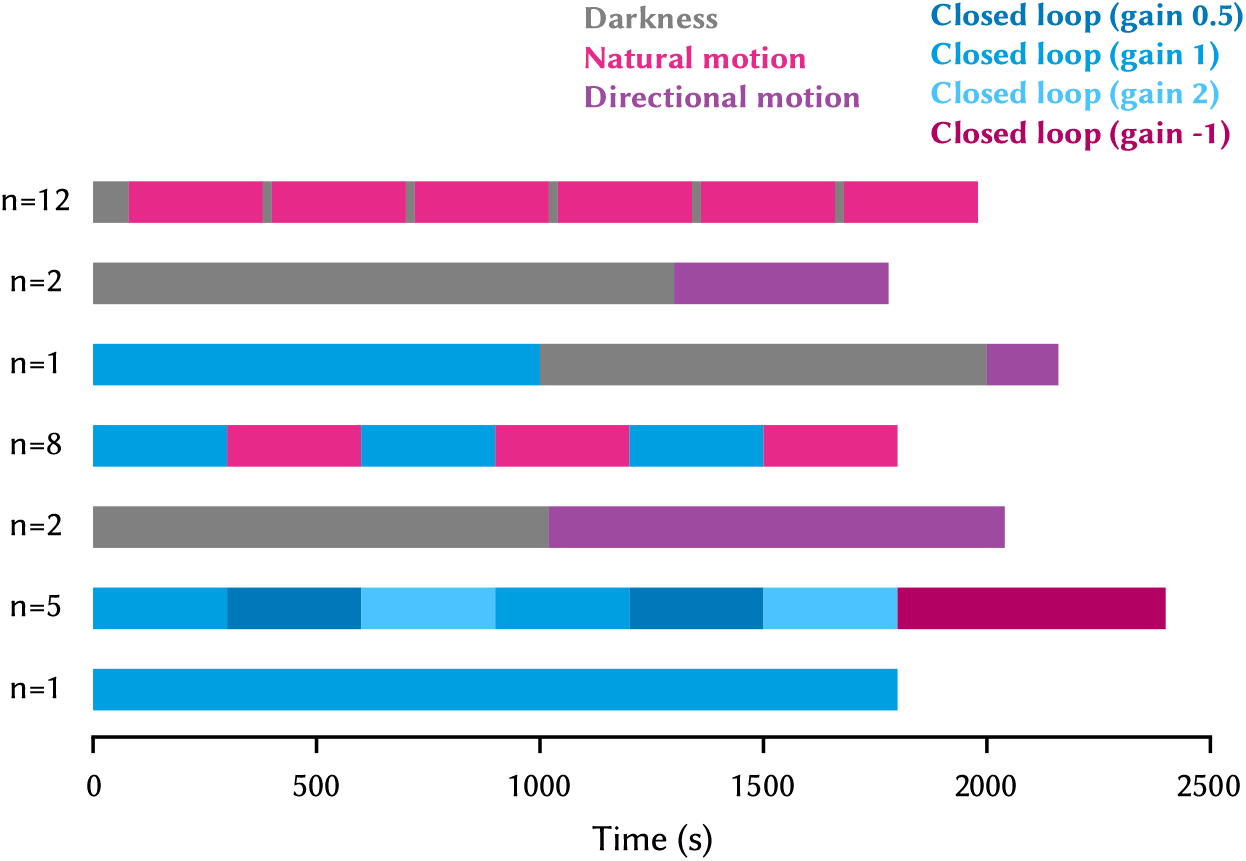
Sequences of visual stimuli presented during all experiments in the dataset.

The paradigm to investigate the role of visual feedback (Figure 12 and Supplementary Figure 9) consisted of an alternation of 5 min of the closed loop and 5 min of the open loop condition (see Figure 25). The paradigm addressing the effect of changing gains (Figure 12 and Supplementary Figure 9) consisted in 5 min blocks of each gain condition, with two repetitions for each condition, in the following sequence of gains: [1, 0.5, 2, 1, 0.5, 2, −1, −1]. The Stytra scripts for the control of the experimental stimuli will be shared together with the rest of the code.

### Data analysis and statistics

All parts of the data analysis were performed using Python 3.7, and Python libraries for scientific computing, in particular `numpy` (Harris et al., 2020), `scipy` (Virtanen et al., 2020) and `scikit-learn` (Pedregosa et al., 2011). The Python environment required to replicate the analysis in the paper can be found in the paper code repository. All figures were produced using `matplotlib` (Hunter, 2007). All statistical tests used were non-parametric, either Mann-Whitney U test for unpaired comparisons (`mannwhitneyu` from `scipy`) or Wilcoxon signed-rank test for paired comparisons (`wilcoxon` from `scipy`). All the analysis code and the source data will be shared upon publication.

#### Lightsheet imaging data preprocessing

The imaging stacks were saved in `hdf5` files and then directly fed into `suite2p`, a Python package for calcium imaging data registration and ROI extraction (Pachitariu et al., 2016). We did not use `suite2p` algorithms for spike deconvolution. As the planes were spaced by roughly 10 μm, we ran the detection on individual planes and did not merge ROIs across planes. Parameters used for registration and source extraction in suite2p can be found in the shared analysis code. The parameter that specifies the threshold over noise that is used to detect ROIs (`threshold_scaling`) was adjusted differently from acquisition to acquisition to compensate for the variability in brightness that we observed from fish to fish. From the raw F traces saved from `suite2p` (`F.npy` file), Δ*F* /*F* was calculated using as baseline the average fluorescence in a rolling window of 900 s, to compensate for some small amount of bleaching that was observed in some acquisition. The signal then was smoothed with a median filter from `scipy` (`medfilt()` from `scipy`), and Z-scored so that all traces were centered on 0 and normalized to a standard deviation of 1. The coordinate of each ROI was taken as the centroid of its voxels. To register all lightsheet experiments to a common coordinate system, we defined manually for each experiment the location over the three axes of a point corresponding on the midline of the fish on the anterior-inferior limit of the aHB, and translate all coordinates so that such point was set to 0.

#### Behavior data preprocessing

The behavioral data was pre-processed to detect swims and extract their properties using the `bouter` package (Štih et al., 2022b). First, the tail trace was processed with a function to reconstruct terminal tail segments that were miss-tracked during the online tracking using an interpolation based on an extrapolation from the reconstructed segments angles and the tail angles at previous timepoints. Then, tail angle was re-computed, and vigor was calculated as the standard deviation of the tail angle trace in a rolling window of 50 ms. Swims were defined as episodes when the vigor crossed a threshold of 0.1 for all fish. For all swims, we then computed the laterality index as the average angle of the tail during the first 70 ms of the swim (Figure 26).

**Figure 26:**
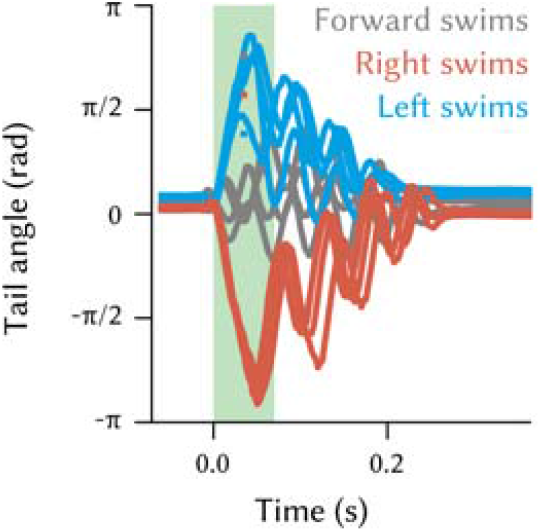
Example left, right, and forward swims, with the window that is used to compute the laterality index.

This value has been shown to correlate well with the angle turned by a fish when swimming freely (Huang et al., 2013; Dragomir et al., 2020). To classify right, left, and forward swims, we fit a trimodal gaussian distribution to the histogram of swim laterality indexes, enforcing the two side curves to be symmetric. Then, we used the intercept of the central and lateral gaussians to determine the threshold used for the swim classification (±0.239 rad).

For eye tracking, the video recording of the entire experiment was processed using the `deeplabcut 2.0` package (Mathis et al., 2018; Nath et al., 2019), a Python pose estimation package based on DeeperCut (Insafutdinov et al., 2016) to detect in every frame four points evenly spaced on each eye. Eye angle was defined as the median angle of the segments that connected the rostral-most point of the eye with all the others. *Gaze direction* was defined as the average of the angles obtained for the two eyes.

#### Detecting r1π neurons

r1π ROIs were first observed to be the ones with the highest anticorrelation with other ROIs in the dataset. Therefore, to detect them, for every experiment we computed the correlation matrix of all traces and selected ROIs that had a correlation below a given threshold with at least another ROI in the dataset. The threshold was manually adjusted for every fish, in order to include as many ROIs that were part of the network as possible, while keeping out other signals. For all fish, the threshold was between −0.75 and −0.5 for both the lightsheet and the two photon experiments. To check that the selected ROIs were convincingly part of the r1π and that we were including enough cells from it, we performed PCA over time using only traces from the selected ROIs and we then looked at the projection of all ROIs onto the first two principal components. When a satisfactory threshold was chosen, most included neurons formed a circular pattern in PC space (see Supplementary Figure 2).

As sometimes some additional ROIs were included, an additional manual step of selection was performed on the correlation matrix of the cells. An optimal sorting of the traces based on their angle in PC space was computed, and the correlation matrix plotted with the same sorting. Then, some traces were excluded based on the amount of discontinuity they would produce in the matrix.

We note that other approaches could be used to parse out those cells, such as restricting the anatomical location where to find them, or including them based on the proximity to some ring fit in PC space. We used just the anticorrelation and exclusion from the correlation matrix to avoid circular reasoning in the observations reported. Future investigations on this system might develop more principled procedures to isolate the r1π neuron population from the rest of the network by the features of their highly constrained dynamics.

With our strategy, we could detect a r1π network in approximately 20% to 30% of the imaged animals. In the rest of the fish, sometimes behavior was just very sparse (a few swims over the entire experiment), or not very directional (only forward swims performed). In other fish, even if behavior was good the anticorrelation criterion could find only a handful of strongly anticorrelated neurons. Although those neurons were likely to be of the described network, as their activity state changed with the occurrence of directional swims, the low number of ROIs made it impossible to properly characterize their population dynamics. Finally, in some fish the rotatory dynamics was observable only in a small temporal interval of the experiment, and they were not included in the dataset.

#### Rotated principal component calculation

We developed a way of registering PC projections from one fish to the other in a way that was consistent with the anatomical distribution of the cells. After computing principal components over time for the r1π neurons, we fit a circle to the projection of the activity of all individual r1π neurons to the first two PCs using the hyper_fit() function from the `circle_- fit` package, a Python implementation of the hyper least square algorithm (Kanatani and Rangarajan, 2011), and rescale and translate the PCs to have a unit radius circle centered on (0, 0).

Then, we computed a weighted average across all the vectors representing ROIs in this two-dimensional space, weighted by their location in the rostro-caudal and the left-right anatomical axes (Figure 27). As a result, we got two vectors, one pointing in the direction of the most rostral ROIs, and the other in the direction of the rightmost ROIs; we then rotated and flipped each fish’s projection so that those two axes matched across fish, *i. e*. the sum of the two angular distances *abs*(*θ*_1_)+ *abs*(*θ*_2_) was minimized (see Figure 27 for the definition of *θ*_1_ and *θ*_2_). We call the axes of this space *rotated principal components (rPCs)*.

**Figure 27:**
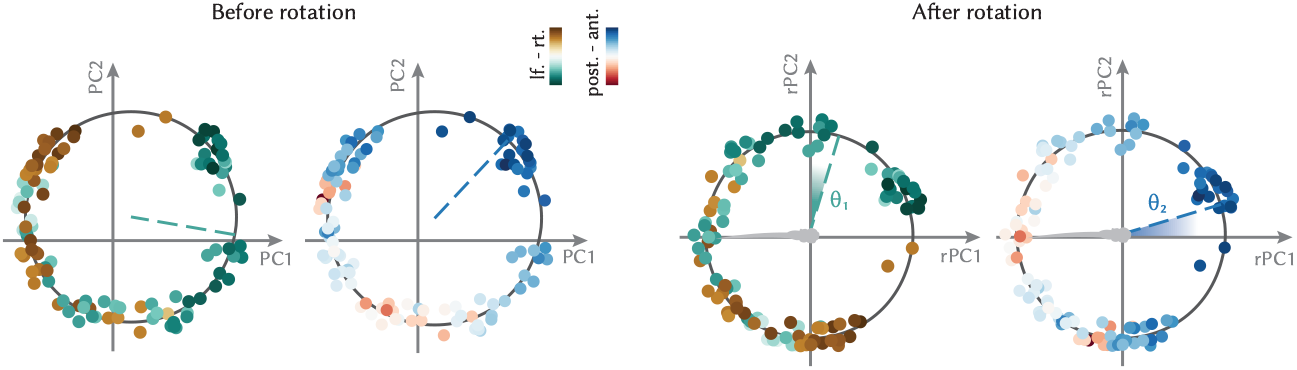
Co-registration of PC projections. *Left*, Projection over the first two PCs calculated over time, color-coded by left-right location of the ROI or anterior-posterior location. The anatomical axes vectors computed by vector average of the ROI projections weighted by their anatomical location are also displayed. *Right*, The projections in rPC space, where the angular distances *θ*_1_ and *θ*_2_ between the anatomical axes and the rPC axes were minimized. The schematics of the fish illustrates the orientation of the rPC projections after the registration.

After having calculated rPCs for an experiment, all ROIs were assigned an angle *α*_*i*_ based on their position over the circle in rPCs space. The convention used for the angle was that:

- *α* ∈ (−*π, π*]
- Caudal neurons had *α* = 0
- *α* increased when moving clockwise in the anatomical location of the neurons

Therefore, looking from above the horizontal plane, leftmost ROIs had *α* = *π* /2, and rightmost ROIs *α* = −*π*/2 (Figure 28).

**Figure 28:**
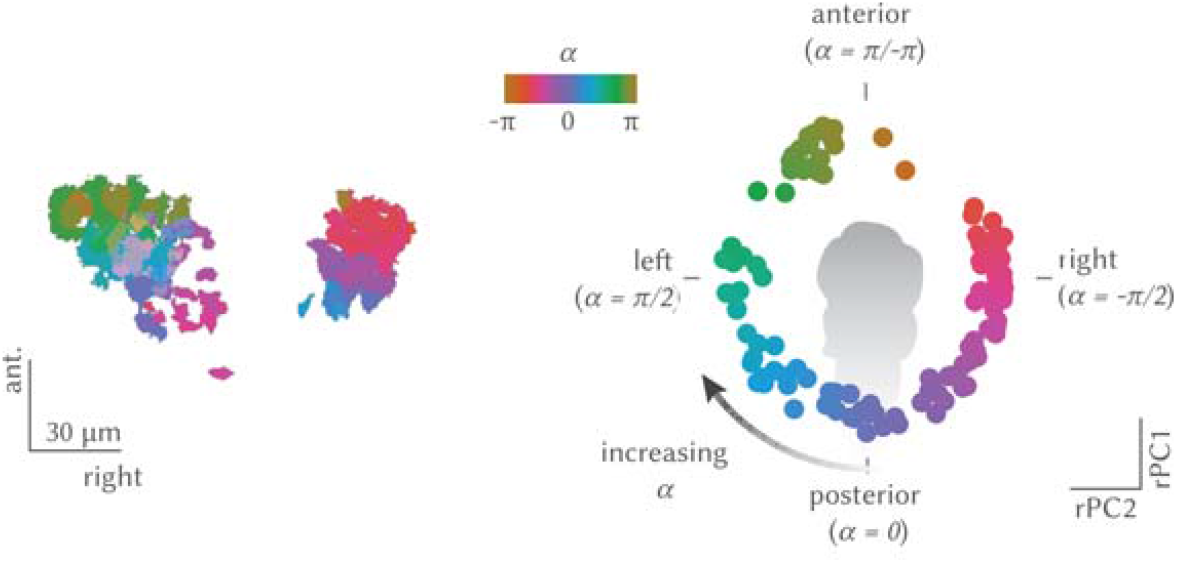
Illustration of the convention used for defining angles in the paper. *Top*: ROIs in rPC space, with labeled *α* and corresponding anatomical position of the ROIs. *Bottom*: the anatomy for the same ROIs, color-coded by *α* in rPC space.

To test the hypothesis that the network is anatomically organized, we used the Fisher-Lee definition of circular correlation coefficient (Fisher and Lee, 1983). We also fit a sinusoidal curve to the distribution of ROIs left-right and anterior-posterior coordinates over the ROIs angle in rPCs space, and compared the fit residuals to the residuals computed over a shuffle computed by reassigning randomly ROI coordinates (Figure 29).

**Figure 29:**
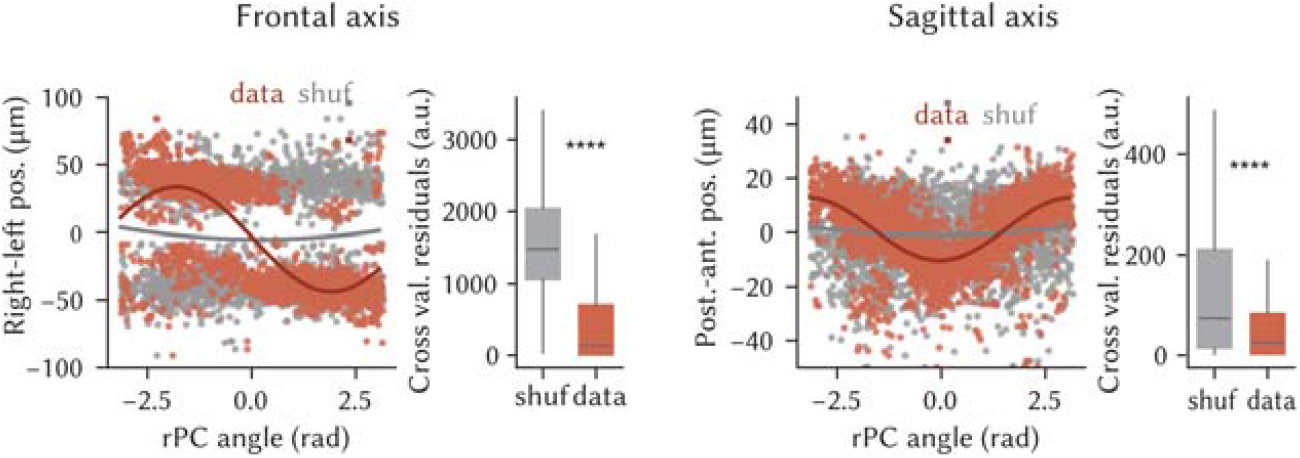
*Left*, Fit of a sinusoidal wave to the anatomical position on the right-left position, as a function of the ROI phase in rPC space, and distribution of square distances of ROIs from the fit (*P* < 0.001, Mann-Whitney U test, n = 1330 ROIs from n= 31 fish). The fit was computed over 50% of the ROIs, and the residuals calculation over the left-out 50%. *Right*, The same, for the antero-posterior axis (*P* < 0.001, Mann-Whitney U test, n = 1330 ROIs from n = 31 fish).

**Figure 30:**
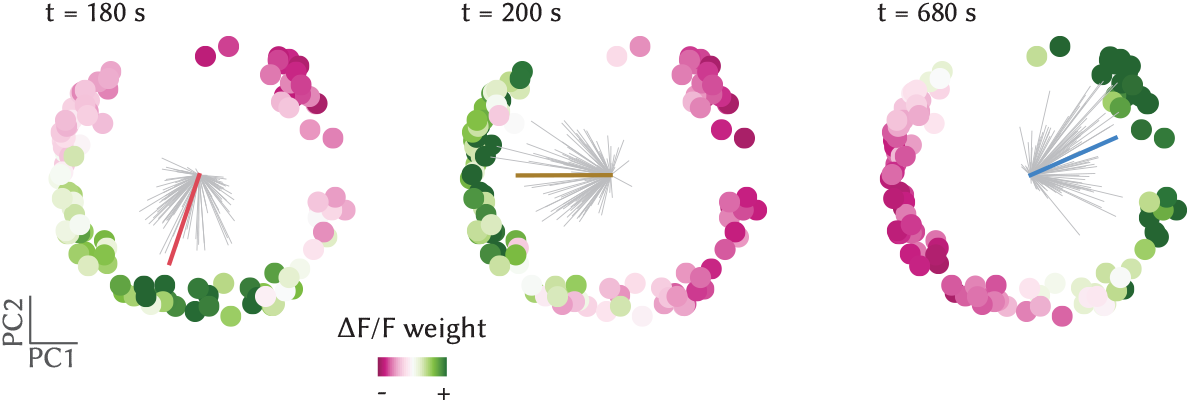
Network phase was computed as the angle of the vector average over all neuron projections in rPC space weighted by their (normalized) activation. Left, center, and right panels show the rPC projections color-coded by the state of activation of neurons at three different time points. Gray lines show the weighted vector of each neuron, and the thick line their average, color-coded by their angle. Note how the angle definition and the colors match the legend defined in Figure 28.

#### Network phase calculation

We derived the phase *ϕ*(*t*) to describe which part of the circle in rPCs space was the most active at every time point (Figure 30, Movie 2). For every frame, we computed a vector average **v** of all the *n* ROI vectors **rPC**_**i**_ in the two-dimensional rPCs space, weighted by the state of activation of each ROI *f*_*i*_ (*t*) (the Δ*F* /*F* at time *t*):

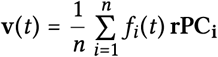

Note that for this vector averaging, the Δ*F* /*F* of all ROIs at time *t* were clipped to their 2% and 98% percentiles and normalized to have mean 0 across ROIs at every time point:

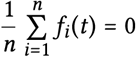

Where **rPC**_**i**_ is the 2-dimensional vector of rPC scores for the *i*^*th*^ neuron, and *f*_*i*_ (*t*) its (normalized) Δ*F* /*F* at time *t*.

The *network phase ϕ*(*t*) is then defined as the angle *ϕ*(*t*) subtended by this vector **v**(*t*) subject to the same conventions as the *α*_*i*_ s defined above Figure 28:

- *ϕ* = 0 corresponds to caudal neurons being active
- increments in *ϕ* correspond to activity rotating clockwise, and decrements of *ϕ* to activity rotating counterclockwise

Therefore, *ϕ* = 0 corresponded to the activation of the network in the rostral part, *ϕ* = *π* /2 to activation of the left part, *ϕ* = ±*π* to activation in the rostral part, and *ϕ* = 2 to activation in the right part (Figure 28).

For all further analyses, the *unwrapped* or cumulative phase was used (unwrap function from `numpy`), *i. e*. every discontinuity at π/-π was removed adding to parts of the trace an offset 2*kπ* for some integer *k*.

#### Calculation of average activity profile

To estimate the average activation profile of the network across the ring of neurons, we started by interpolating the neuron’s traces to a matrix spanning the interval −*π* to *π* in 100 bins (Figure 31).

**Figure 31:**
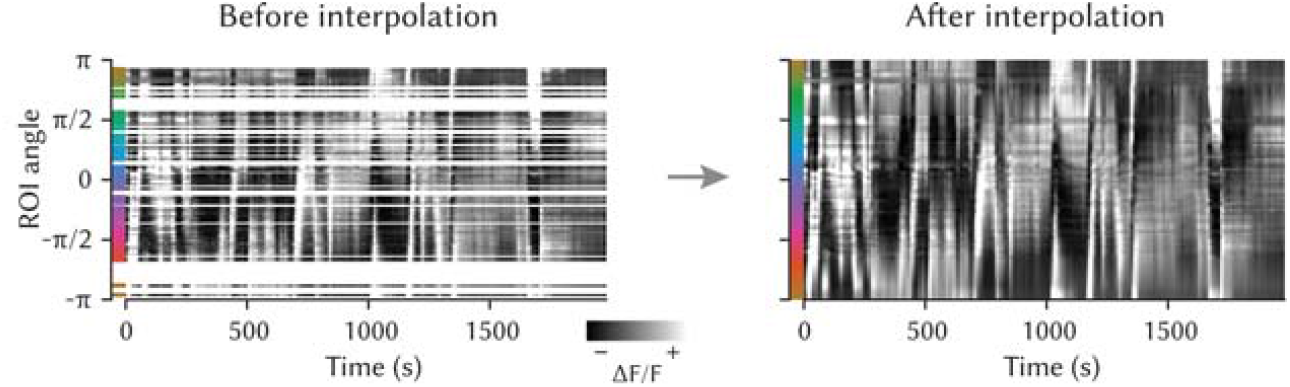
Interpolation of network activity from neuron angles. *Left*, Traces of individual neurons sorted and spaced in *y* using their angle *α*_*i*_. The colors on the left map neuron angles. *Right*, The same activity, after interpolating the activation between −*π* and *π*.

Then, we circularly shifted each column of the matrix so that the phase, and hence the network activation peak, was always positioned at the center of the matrix (Figure 32). Finally, we calculated the average and standard deviation of the matrix across the time axis. To make sure the result was not the consequence of the resampling procedure, we also performed the circular shift of the raw matrix of traces, sorted according to neurons’ *α*_*i*_, and we got consistent results (Supplementary Figure 6a,b).

**Figure 32:**
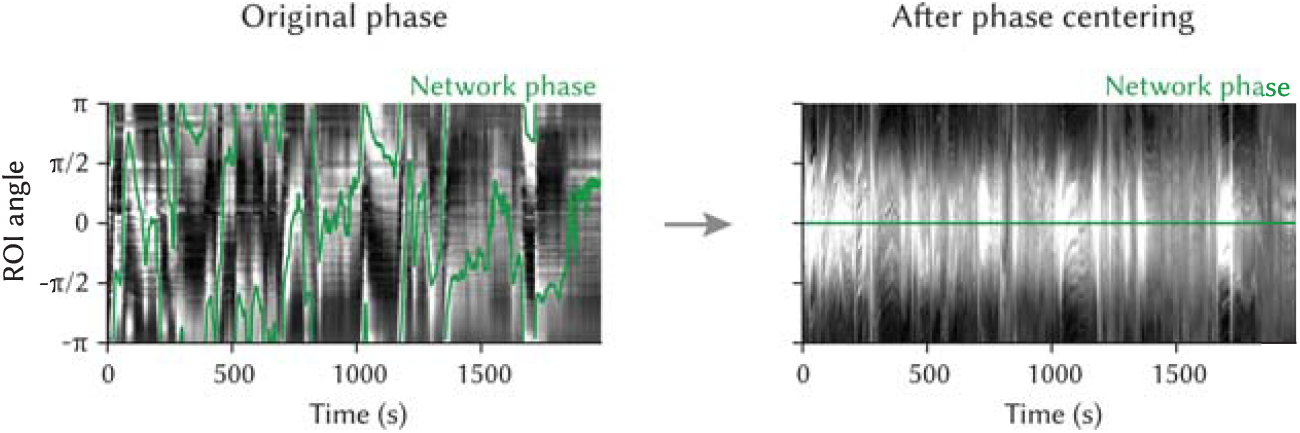
Phase-zeroing process: for every time point, a circular permutation of the (interpolated) activity matrix was computed so that the peak of activation, mapped by the phase (*left*), was always in the center of the matrix. *Right*, The matrix of traces, after the interpolation.

**Figure 33:**
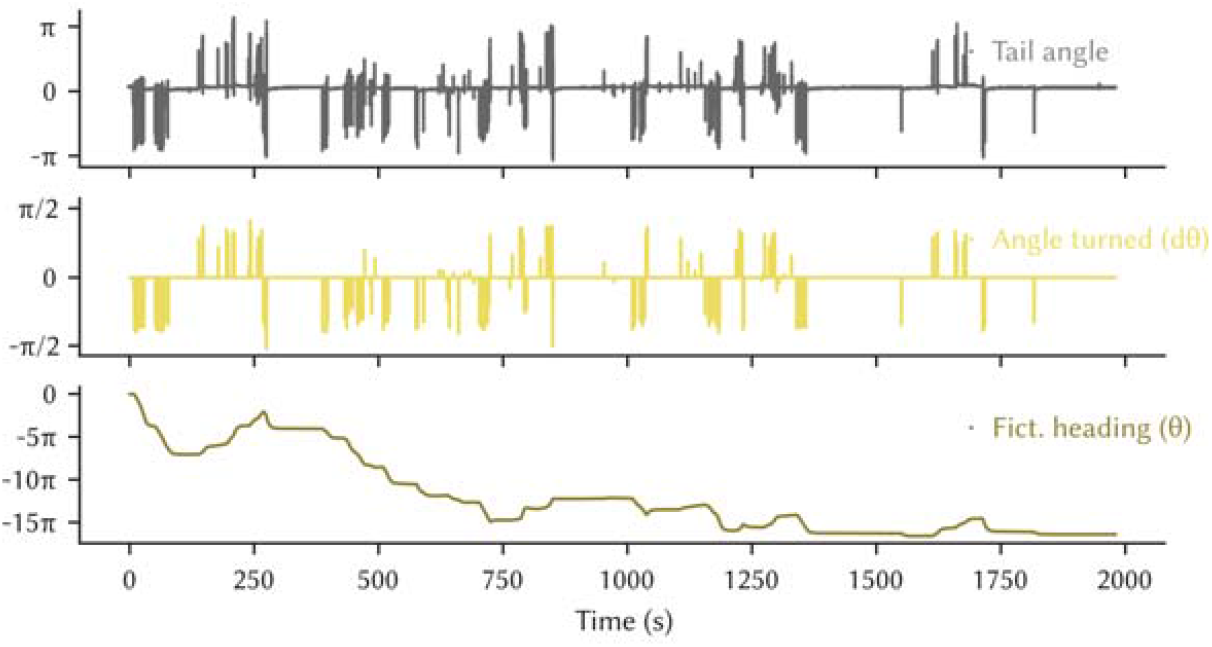
Calculation of the fish heading estimate. *Top*, Raw tail trace. *Middle*, Laterality index for each swim. *bottom*, Cumulative sum of laterality indexes of swims.

#### Estimated heading calculation and correlation with phase

To compute estimated heading for the analysis reported in Figures 10 and 11, we estimated the instantaneous angular velocity as the laterality index value for each individual swim (see *Behavior data preprocessing*), and we integrated it over time to obtain an estimated heading direction for the head-restrained fish.

We note that although the relationship between the laterality index and the fish orientation change in freely swimming animals is highly linear, the slope of the linearity is not necessarily one; moreover, the precise extent of the tail that is tracked, the embedding procedure, and the fact that the head is immobilized in agarose for our head-restrained imaging experiments are all parameters that can impact on the precise kinematic of the tail movements and make a precise numerical comparison between head-restrained and freely swimming experiments difficult. Therefore, we did not aim at reconstructing a fully realistic estimated heading direction, and relied on quantifications that either captured just the correlation between estimated heading changes and network phase changes, or quantified the slope coefficient between the two quantities in relative comparisons within one experiment (for the visual feedback and gain change experiments).

For the results reported in Figure 11, we calculated for each fish the correlation between heading and phase in a rolling window of 300 s (10 overlaps for each window), and the same correlation but using a non-overlapping 5 min epoch of the heading trace for the shuffle distribution. The moments reported in Figure 11 refer to this population of intervals and shuffle intervals for each fish.

#### Swim-triggered and saccade-triggered analyses

For the directional swim-triggered and saccade triggered analysis of Figure 8 and Supplementary Figure 11, we cropped for each fish the phase around each event, we computed a fish average for all curves with at least 3 cropped samples, and we subtracted the mean of the 10 s interval before the event.

#### Heading/phase slope fitting for visual feedback experiments

In the experiments reported in *The r1π network is not affected by visual inputs*, we wanted to quantify whether the presence of closed-loop visual feedback or the effect of different gain parameters of the closed loop visual feedback had an effect on the relationship between the change in heading and the phase of the network. As swims often happen in sequences and the average network phase change seems to plateau after approx. 5s from the focal swim, we decided to look at the relationship between the amount of phase changed in a window between 15 s to 20 s after the swim, and the amount of estimated heading change in the same interval (which will potentially accumulate also the effect of other swims in the sequence).

The choice of the window was arbitrary, and all the results hold with other intervals in the 5s to 20 s range. To quantify this relationship, we performed linear regression on the (Δ_*heading*_, Δ_*Phase*_) pairs for all swims in each experimental condition (Figure 34 shows this calculation for all fish), and we compared the values of the regression slope across conditions (Figure 12).

**Figure 34:**
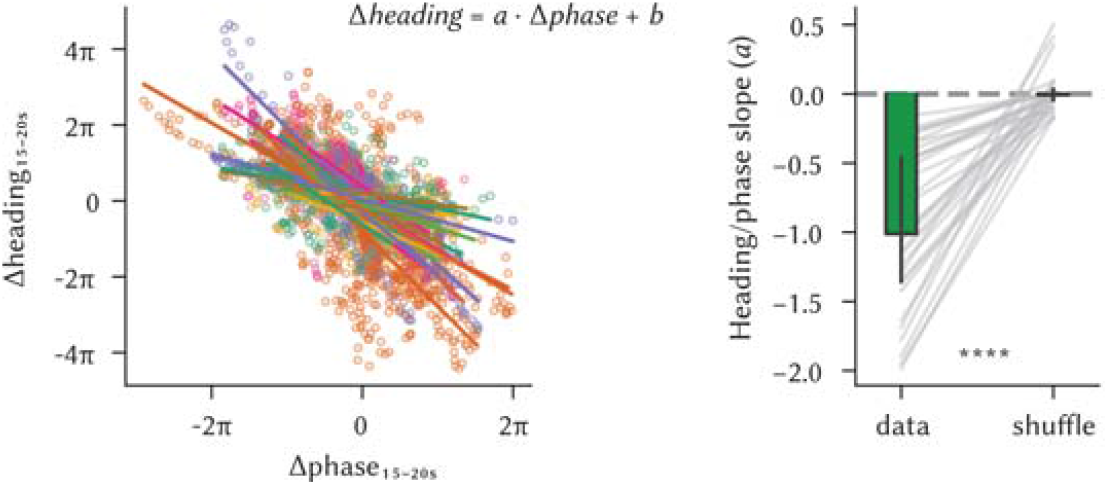
To quantify the effect of visual feedback on the network activity, we performed a linear regression between the amount of shift in the network phase and in the estimated heading, between 15 s after each swim (see Materials and Methods). *Left*, Scatter plot of heading change vs phase change after every swim (individual points, color-coded by fish), and the linear fit for every fish. *Right*, Comparison between the slopes in the data vs. a shuffle over swim identity (data: (median = −1.01, Q1 = −1.36, Q3 = −0.437, n = 31 fish); shuffle: (median = −0.006 47, Q1 = −0.0452, Q3 = 0.0343, n = 31 fish); *P* < 0.0001, Wilcoxon test).

**Figure 35:**
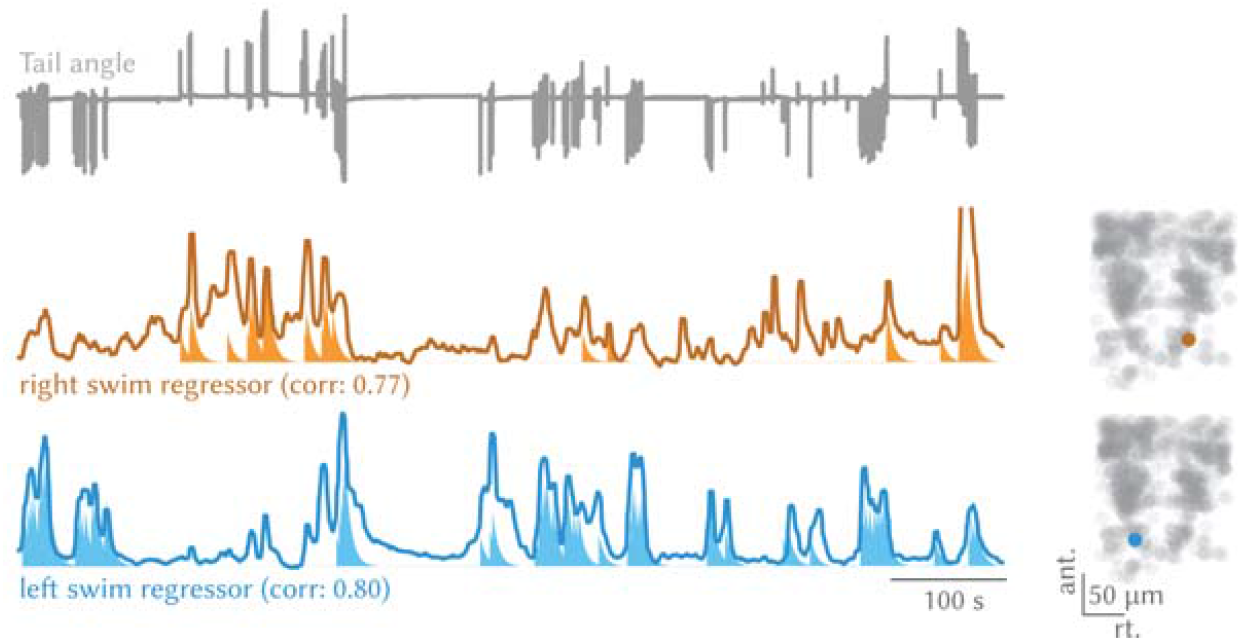
*Left*, Example traces for right and left swims-selective neurons together with the fish tail trace, and *right*, their anatomical location, shown on top of a scatter plot of all ROIs from the same fish. The regressors are shaded below the traces, and the correlations of the traces with the regressors are reported in the plot.

**Figure 36:**
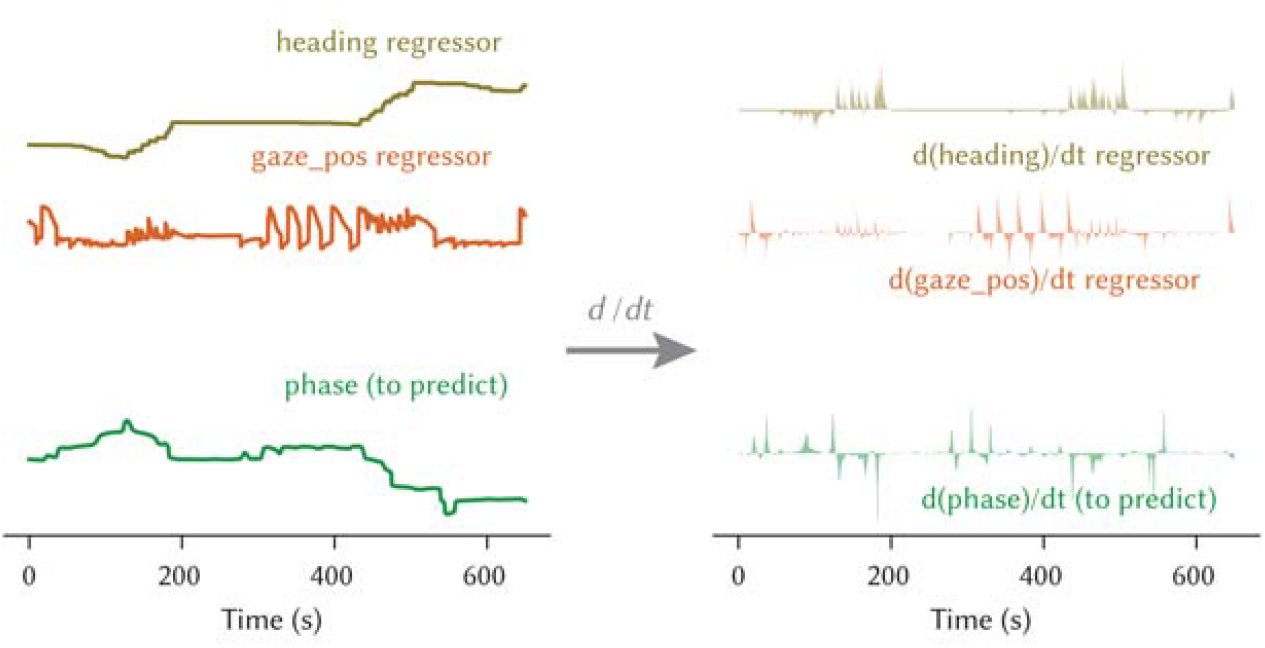
Illustration of the procedure for fitting heading and gaze derived regressors to the phase (*left*). Linear regression or multiple linear regression was performed on the time derivatives of those quantities (*right*).

#### Left and right swim and gaze angle regression

To understand whether in the region there was activity related to left and right swims, we performed a regressor-based analysis. A set of regressors was built by convolving with an exponential decay function an array that was zero everywhere and 1 in correspondence of either left or right swims (for Supplementary Figure 10) or with the gaze direction array (for Supplementary Figure 11).

The time constant used was 3 s; the value was higher than the GCaMP6s time constant, but was chosen as it matched more closely the experimentally observed curves. The exact value of the time constant was not critical for the reported results. Each cell’s fluorescence trace was then correlated with both regressors, and the correlation values were used for the analysis and visualizations in Figure 13 Supplementary Figure 10 and Supplementary Figure 11b,c. In the maps of Figure 13 and Supplementary Figure 10b,c left- and right-swim related cells were defined by including in the category cells with a correlation with left- or right-swim regressor > 0.7, and correlation with the other regressor < 0.7.

#### Multilinear regression of eye and tail to network phase

For addressing the relationship between network phase and eye motion, we used the gaze direction, computed as the average between the two eyes angles. For the regression analysis, we used the gaze velocity or the instantaneous fish angular velocity estimated from the swim laterality indexes (both convolved using the same tau as in *Left and right swim and gaze angle regression*), either alone or in combination to fit the temporal derivative of the (unwrapped) network phase.

As a multilinear regression is likely to outperform the regression using only one of the two regressors just by overfitting, we cross-validated the analysis by first calculating the regression values on a randomly drawn epoch of 5 min of the experiment, and calculated the correlation of the phase derivative and the predicted phase derivative in a test 5 min epoch, drawn randomly by making sure it did not overlap with the fit window. The random sampling was repeated 500 times, and the plot in Supplementary Figure 11 reports for each fish the moments of the population of these draws.

#### SBEM data skeletonization

The first reconstructions of cells in the aHB with processes in the IPN were observed by seeding for reconstruction dendrites or axons in the IPN and reaching from there somas in the aHB, in the context of a (still unpublished) broader reconstruction effort. The IPN location in the SBEM stack was firstly inferred by the recognizable organization of the neuropil and cell somata in the rhombomere 1 ventral region. Then, it was confirmed by the tracing of axons that could be reconstructed back to the habenulae through a long bundle of fibers unambiguously identifiable as the fasciculus retroflexus by its course (unpublished data). After those first observations, additional cells with the soma in the aHB were seeded based on the similarity of their processes with already reconstructed cells.

Skeletonization was performed manually by a team of annotators at ariadne.ai ag (Buchrain, Switzerland). Annotators were instructed to flag difficult locations without extending the skeleton at those locations, and to stop tracing after a total time of 2h was reached. At that point, or when a cell was completed, a quality check was performed by an expert annotator. Difficult locations were then decided by the expert, and sent back to the annotator team for additional tracing if necessary. This procedure was iterated until all cells were fully traced. The skeletons were then annotated to distinguish the dendrite and the axon by their morphological features (processes thickness and presence of presynaptic boutons) independently by Ariadne expert annotators or the authors, with convergent results. All further analyses and quantification of the reconstructions were performed using Python. To calculate the centroid position of dendrite and axon for the analyses in Figure 18, we took the average coordinate of the coordinates (in IPN reference space) of all the branching points of dendrites and axons. To generate the distance plot in Figure 20, bottom we calculated for every branching point of every neuron the distance along the frontal and sagittal axis of all the other branching points (of both axons and dendrites) and show the distribution of such distances.

#### Anatomical registrations

To work with the anatomical spaces and their annotations, we used the BrainGlobe `bg-atlasapi` package (Claudi et al., 2020) and either the larval zebrafish brain reference MapZeBrain (Kunst et al., 2019), or a custom lab reference of the aHB and IPN region that will be published together with the data, created by morphing together stacks from different lines using either `dipy` (Garyfallidis et al., 2014) or `CMTK` (Rohlfing and Maurer, 2003). To visualize functional data in the references, an average anatomy computed after centering all stacks with the centering point described in (*Lightsheet imaging data preprocessing*), and then a manual affine registration was performed to the IPN reference. A similar procedure was used to map the electron microscopy data. From the skeletons, a density stack was computed in which the shape and features of the IPN were prominently visible. An affine matrix transformation was found to match this stack on the IPN reference, and used for transforming the neuron’s coordinates. The masks delimiting the IPN and the dIPN were drawn in the IPN reference atlas looking at the localization of habenular axons afferents to the region.

#### 2d auto-correlation of the neuronal activity

For the plots reported in Figures 19 and 20, two-photon microscopy images from a single plane in the IPN were aligned to the frontal and sagittal axes of the brain. The dorsal IPN in the images were masked by manual drawing. The area inside of the mask was divided into 3.5 μm× 3.5 μm square bins. The average fluorescence signal at each bin was z-scored. For each bin, Pearson correlation of the signal traces between the focal bin and all other bins were computed (as shown in Figure 19), and sorted in two dimensions by the distances between two bins in the frontal and sagittal axes (for Figure 20 and Supplementary Figure 16). The correlation coefficients at the same distance were averaged across bins for each animal, and then averaged across animals.

## Data availability and code availability

All the source data used in the functional imaging analysis (raw Δ*F* /*F* traces, ROI maps/coordinates, behavioral traces, and stimulus log from Stytra) and for the anatomical observations (confocal/two-photon stacks, SBEM skeletons) will be deposited immediately upon publication. Similarly, all the scripts for the stimuli generation, data preprocessing, analysis and figure generation will be deposited immediately upon publication.

## Acknowledgements

We thank Winfried Denk for sharing the electron microscopy dataset of a 5 dpf larval zebrafish brain used in this manuscript. We thank Andreas Kist for generating the Tg(elavl3:H2B-mCherry) line. We thank Herwig Baier for sharing the *Tg(gad1b/GAD67:Gal4-VP16)mpn155, Tg(UAS:GCaMP6s)mpn101*, and *Tg(UAS:Dendra-kras)s1998t* lines. LP would like to thank Tommaso James Grossi for insightful discussions on data analysis. The authors thank Shuhong Huang, Virginia Palieri, Emanuele Paoli and the Portugues Lab for discussions on data interpretation. This work was supported by grants to RP from the Volkswagen Stiftung and by the DFG under Germany’s Excellence Strategy within the framework of the Munich Cluster for Systems Neurology (EXC 2145 SyNergy – ID 390857198).

## Author Contributions

LP, HL and RP conceived the project and designed the experiments with help from VS based on preliminary observations from LP. LP performed lightsheet experiments with help from HL and analyzed the data. HL performed anatomical and functional two-photon experiments and analyzed the data, all with help from YKW. FS acquired electron microscopy data and supervised the tracing and LP selected the neurons and analyzed the morphologies. LP, HL and RP wrote the manuscript, with help from YKW.

## Supplementary Figures

**Supplementary Figure 1:**
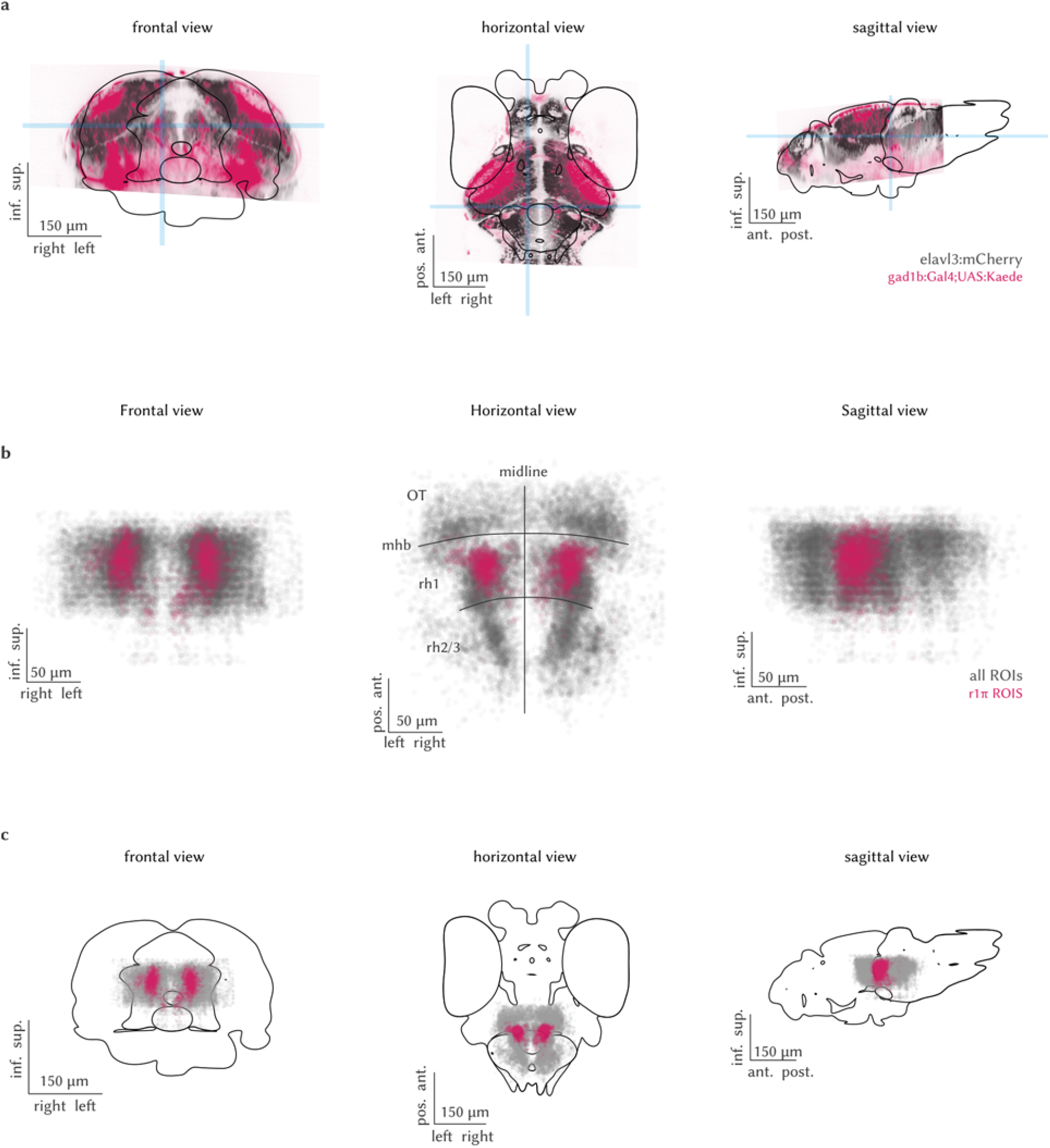
Anatomical location of r1π neurons. a, Frontal, horizontal and sagittal projection of the expression pattern of the *gad1b:Gal4* used in the imaging experiments from one fish. In the gray background, the expression pattern of *elavl3:H2B-mCherry*, on a second channel in the same fish. The blue shades indicate the slices of the stack that were averaged to obtain the views, and are centered on the location of the imaged GABAergic nuclei in the aHB. b, The same views for the r1π neurons in the imaging experiments registered in a common anatomical space (pink), visualized together with all the ROIs extracted from the same experiments. OT: optic tectum, mhb: midbrain/hindbrain boundary, rh: rhombomere. c, The same views for coordinates shown in (b), now registered on the mapzebrain atlas.

**Supplementary Figure 2:**
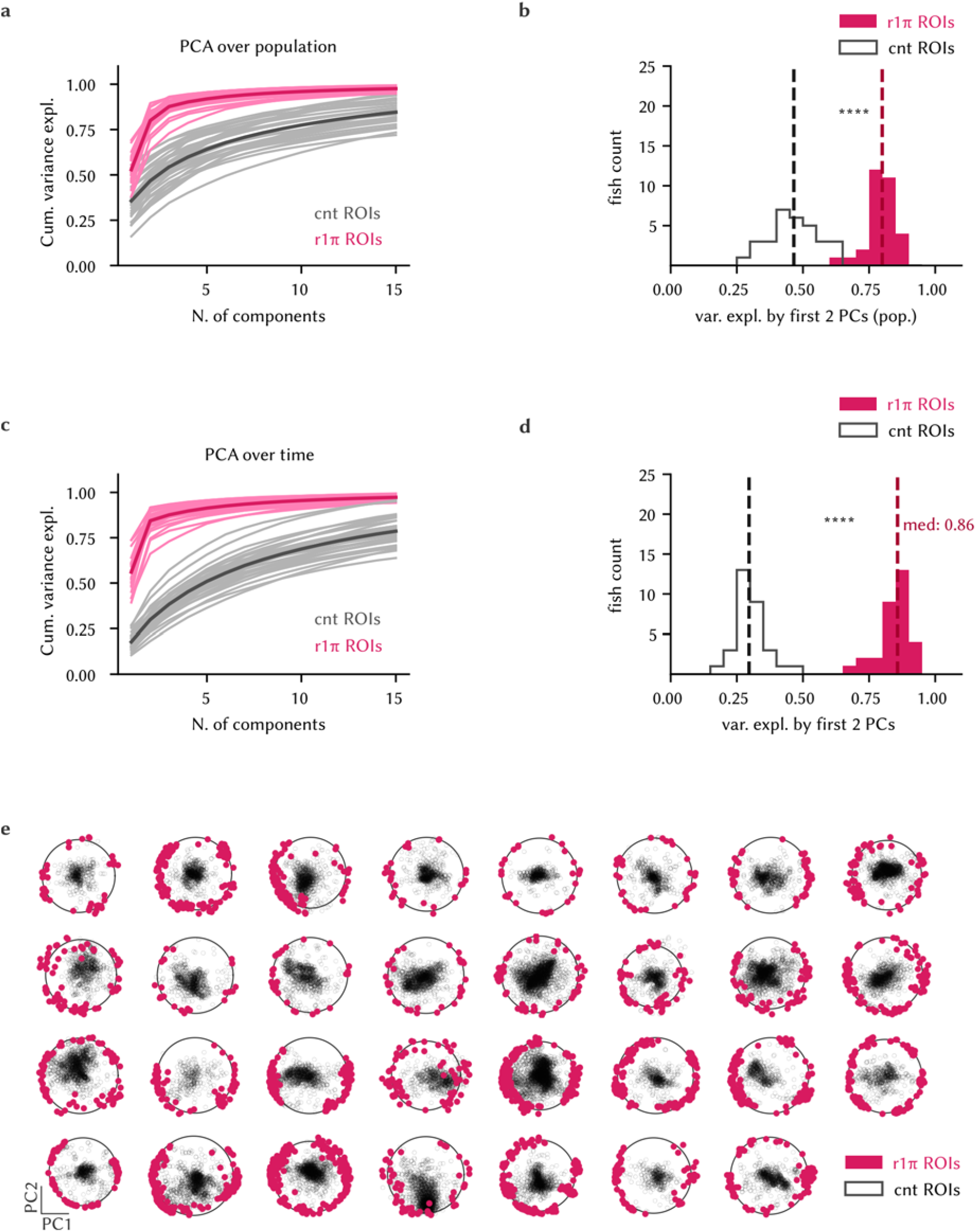
PCA decomposition of r1π neurons activity. a, Cumulative relative variance explained by the first 15 principal compo-nents from PCA decomposition over population for r1π neurons and a population of randomly drawn neurons from the same imaging experiments matching in number the r1π neurons. Light lines: individual fish, dark thick line: population average. b, Variance explained by the first two PCs in the plot in (a), compared between r1π and control neurons (Wilcoxon test, *P* < 0.0001). c, Cumulative variance explained by the first 15 principal components of PCA decomposition over the time dimension, legend as in (a). d, Variance explained by the first two PCs in the plot in (c), compared between r1π and control neurons (Wilcoxon test, *P* < 0.0001). e, Projections over first two principal components calculated over the r1π neurons for all neurons of each fish, for all fish in the dataset.

**Supplementary Figure 3:**
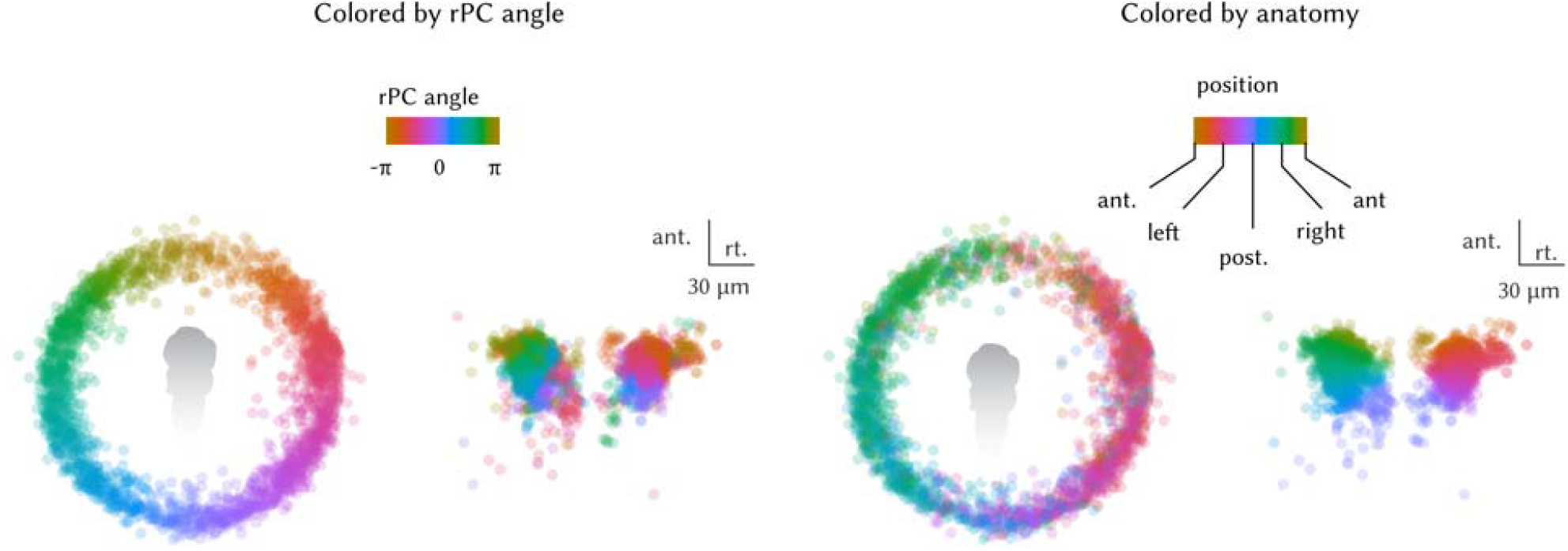
Anatomical organization of r1π neurons angles. *Left*, Projection of ROIs pooled from all neurons in the registered rPC space, color coded by angle in rPC space and their anatomical distribution. *Right*, The same plots, now color coding the angle in anatomical space.

**Supplementary Figure 4:**
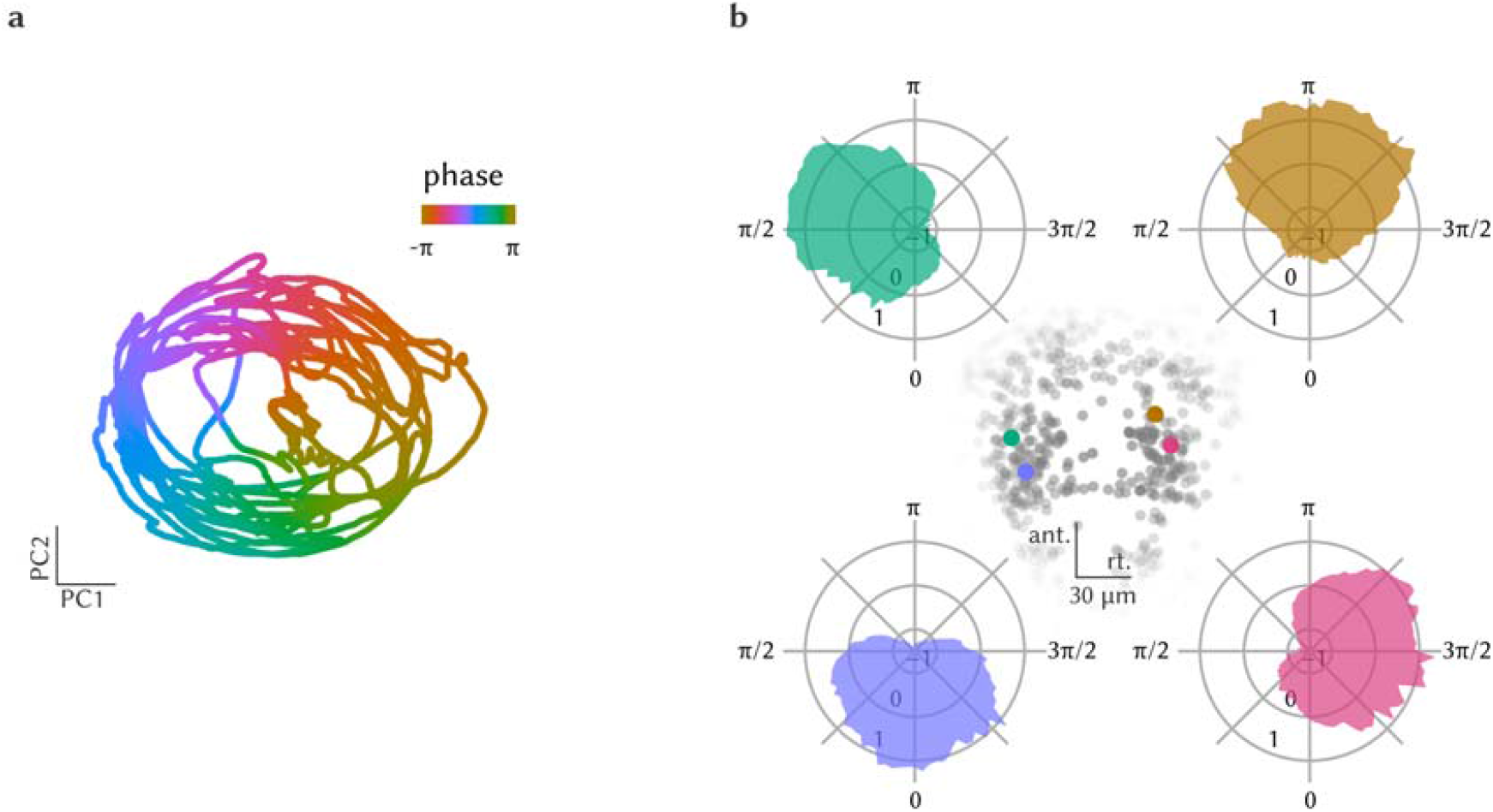
Network phase calculation. a, Trajectory of the network in PCA-reduced phase space, color-coded by the network phase. b, Polar plots showing tuning curves of individual neuron activations as a function of network phase from one fish. Each panel shows the curve for a neuron, color coded by their angle *α*. The anatomical locations of the four neurons are represented in the central inset.

**Supplementary Figure 5:**
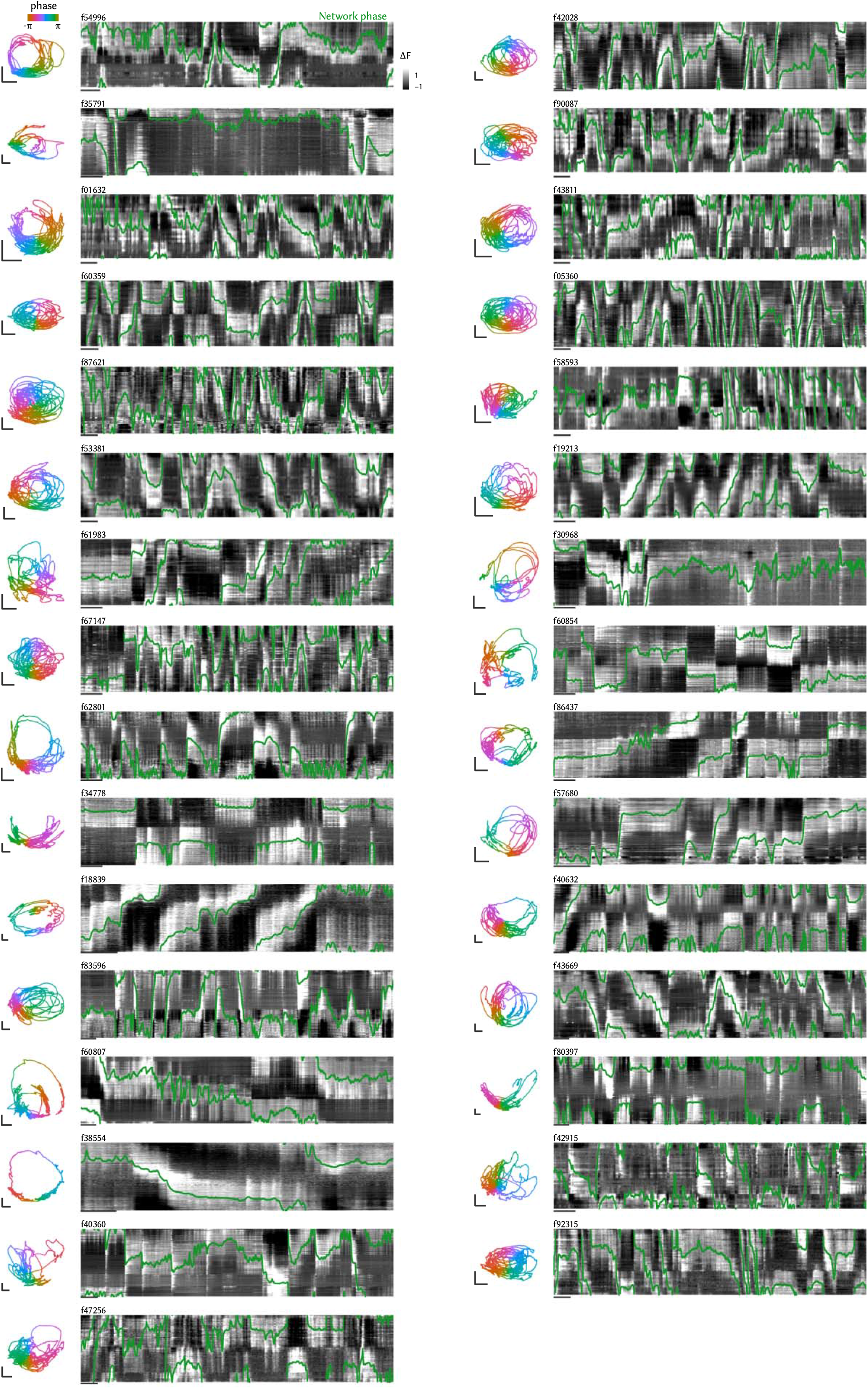
Summary plot for all fish. Raw data showing for each fish of the dataset the trajectory of the network in PC space over the entire duration of the experiment color-coded by network phase (panel on the left), and the raw traces of r1π neurons sorted by neuron angle *α* and network phase in green. The scale bar in the phase space has length of 5, and the bar below the traces indicates 100 s.

**Supplementary Figure 6:**
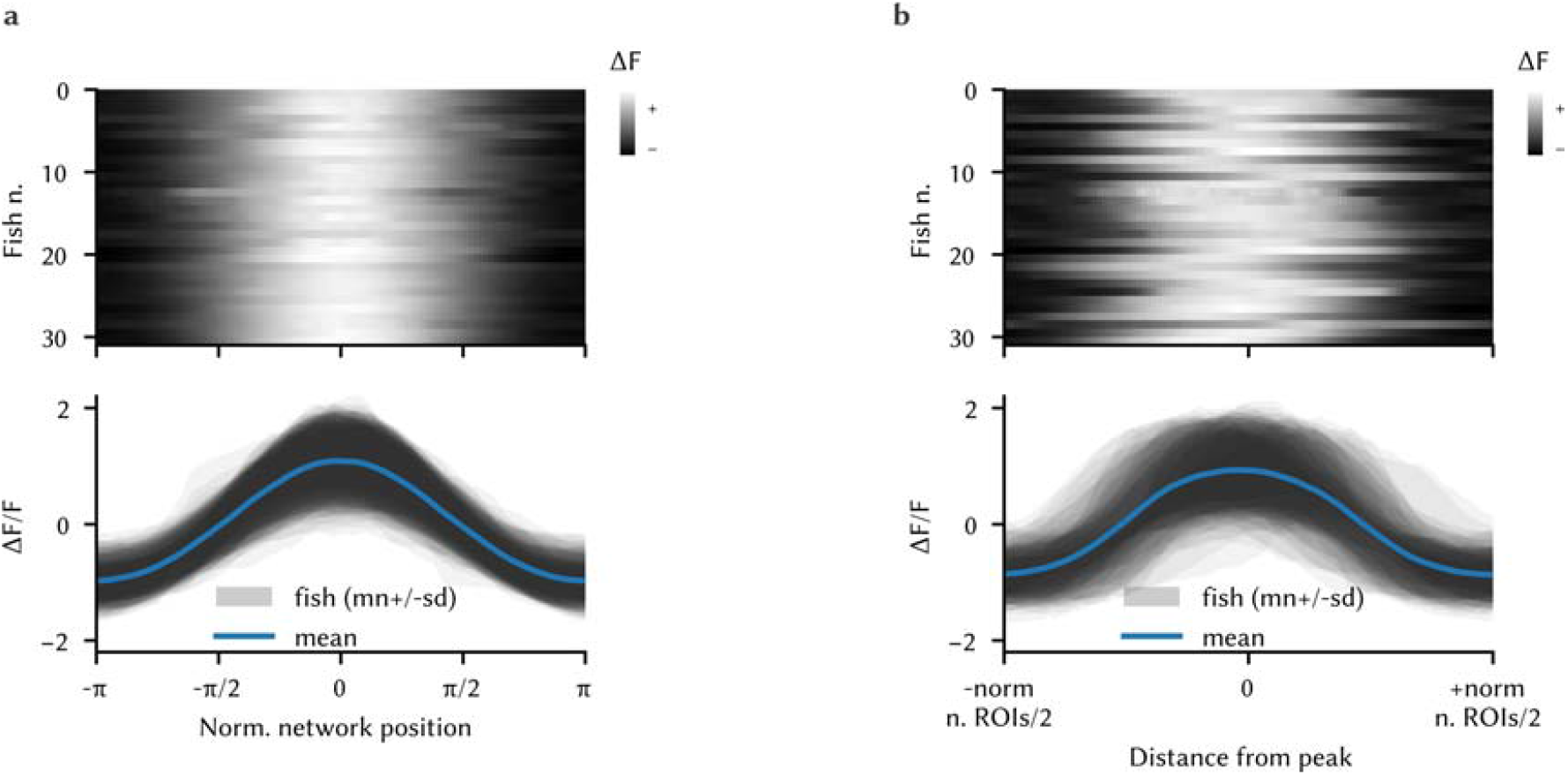
Network activity profile. a, Average activation profile for all fish in the dataset (n = 31 fish). *Top*, Matrix showing the average activation profile for all fish in the dataset and *bottom*, mean ±std over time for each fish (shaded areas) and population average. b, Same plots of panel (c), computed by phase-zeroing in the traces matrix without interpolation.

**Supplementary Figure 7:**
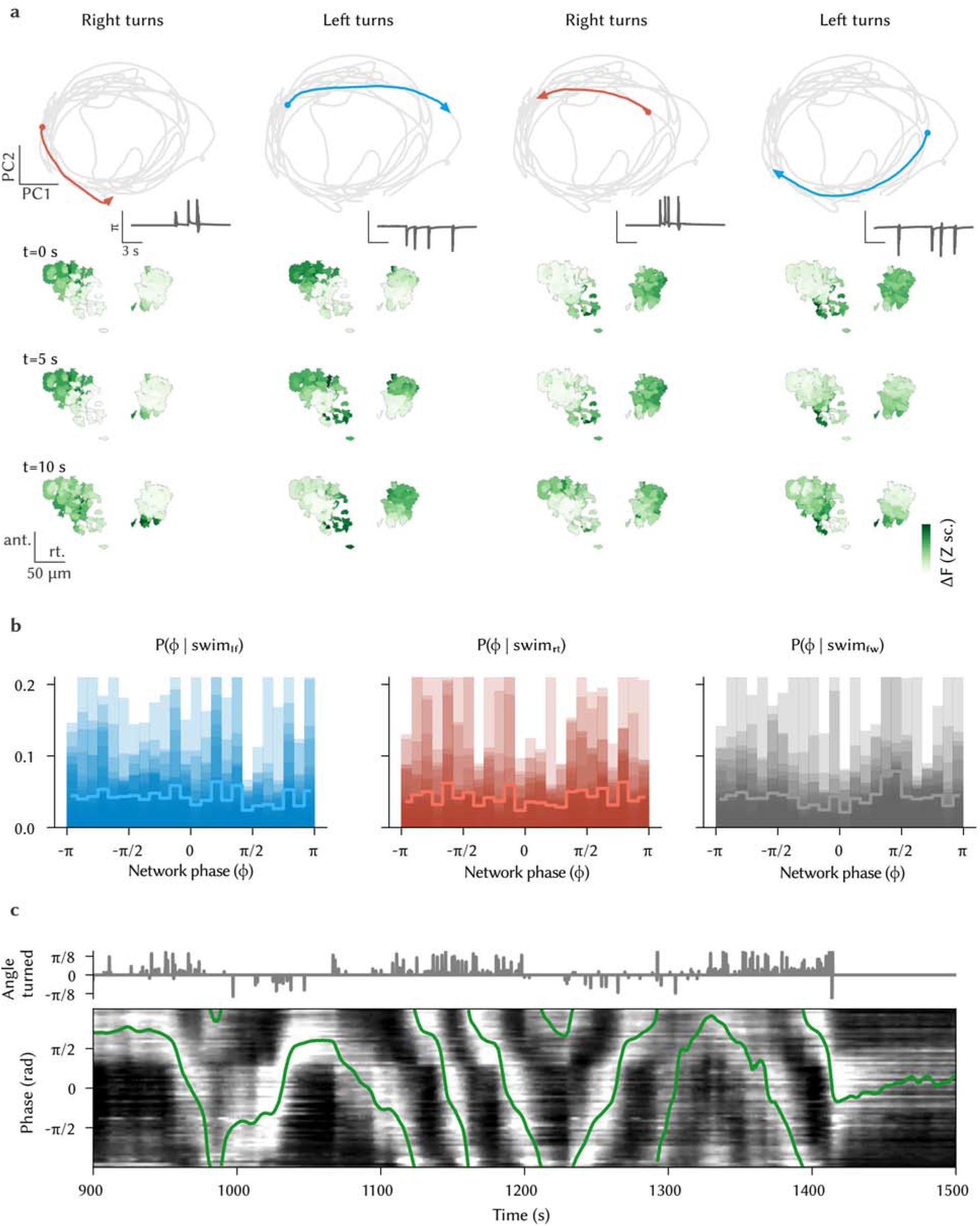
Phase dynamics during directional swims. a, Network trajectory during sequences of left and right swims (same as Figure 7, with more examples). *Top*, trajectory in phase space during a sequence of left swims (see tail angle in the insert). *Bottom*, state of activation of the network before and after the sequence. The four columns show four sequences of left and right turning, with the network starting at different phases. b, Probability of network phase given that a forward, left, or right swim occurred (shaded areas: individual fish; line: population average, n = 31 fish). The distribution is flat, suggesting that the network phase is not instructive with respect to the direction of swimming. c, Example of clockwise and clockwise shifts traversing multiple times the entire network during sequences of repeated directional swims.

**Supplementary Figure 8:**
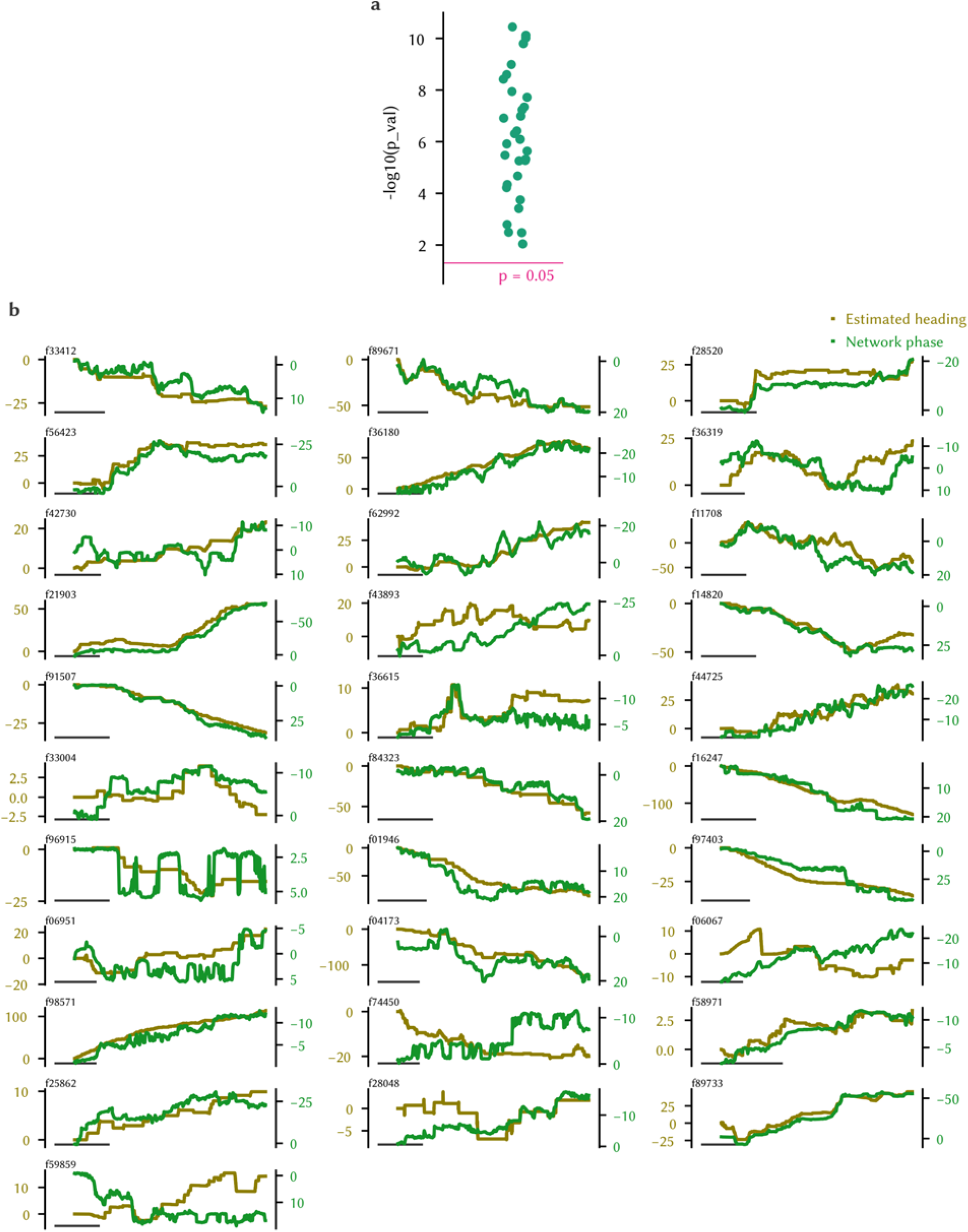
Phase predicts fictive heading. a, Distribution of *P* values from the comparison of correlation of phase and heading in chunks of 5 min in the data and a shuffle (Wilcoxon test, < 0.01 for all fish). b, Raw estimated heading and phase for all fish in the dataset.

**Supplementary Figure 9:**
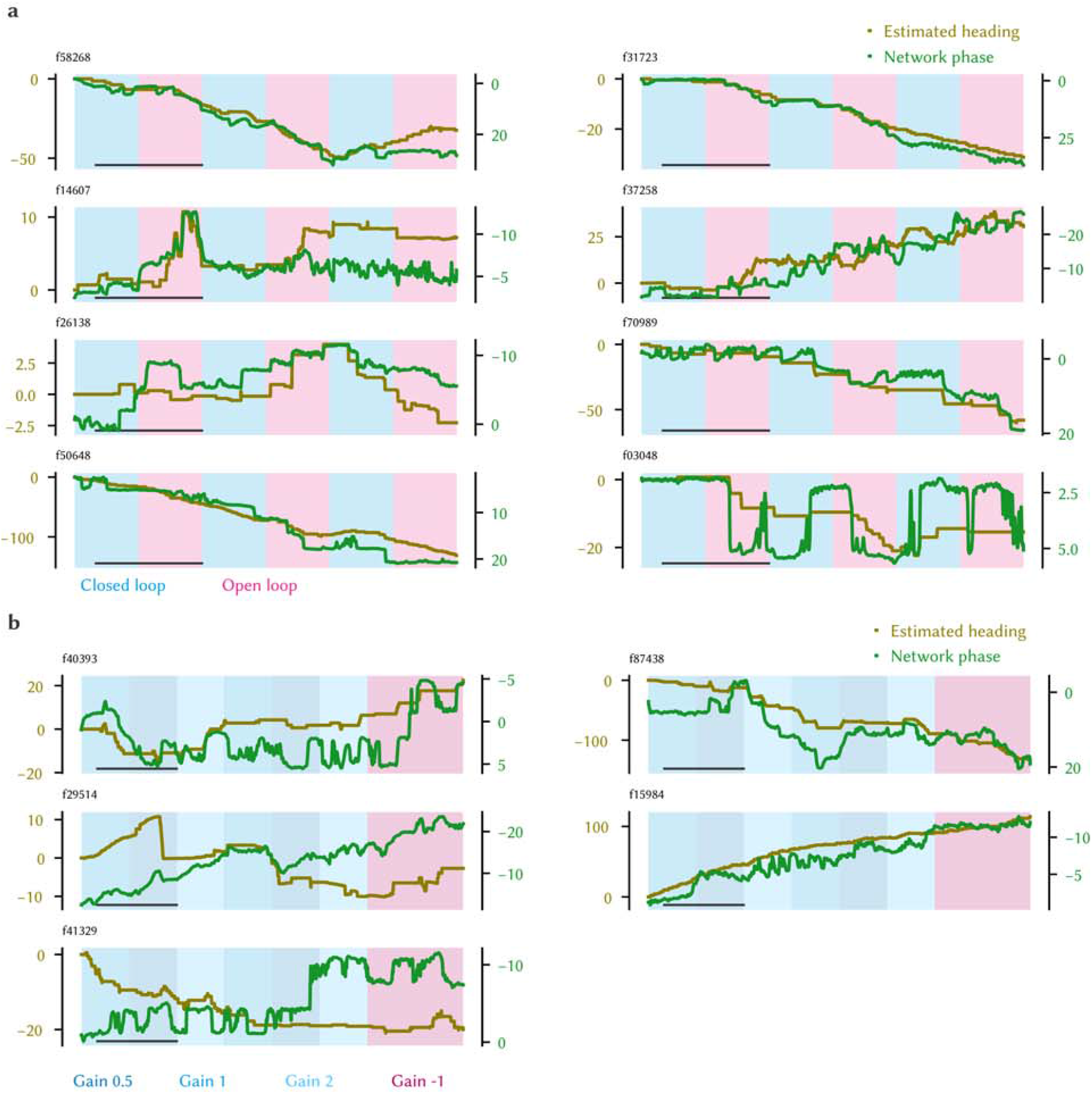
Visual feedback is not required for the turning integration a, Heading, phase, and the experiment condition for the closed-loop/open-loop experiments (scale bar: 500 s). b, Heading, phase, and the experiment condition for the gain modulation experiments.

**Supplementary Figure 10:**
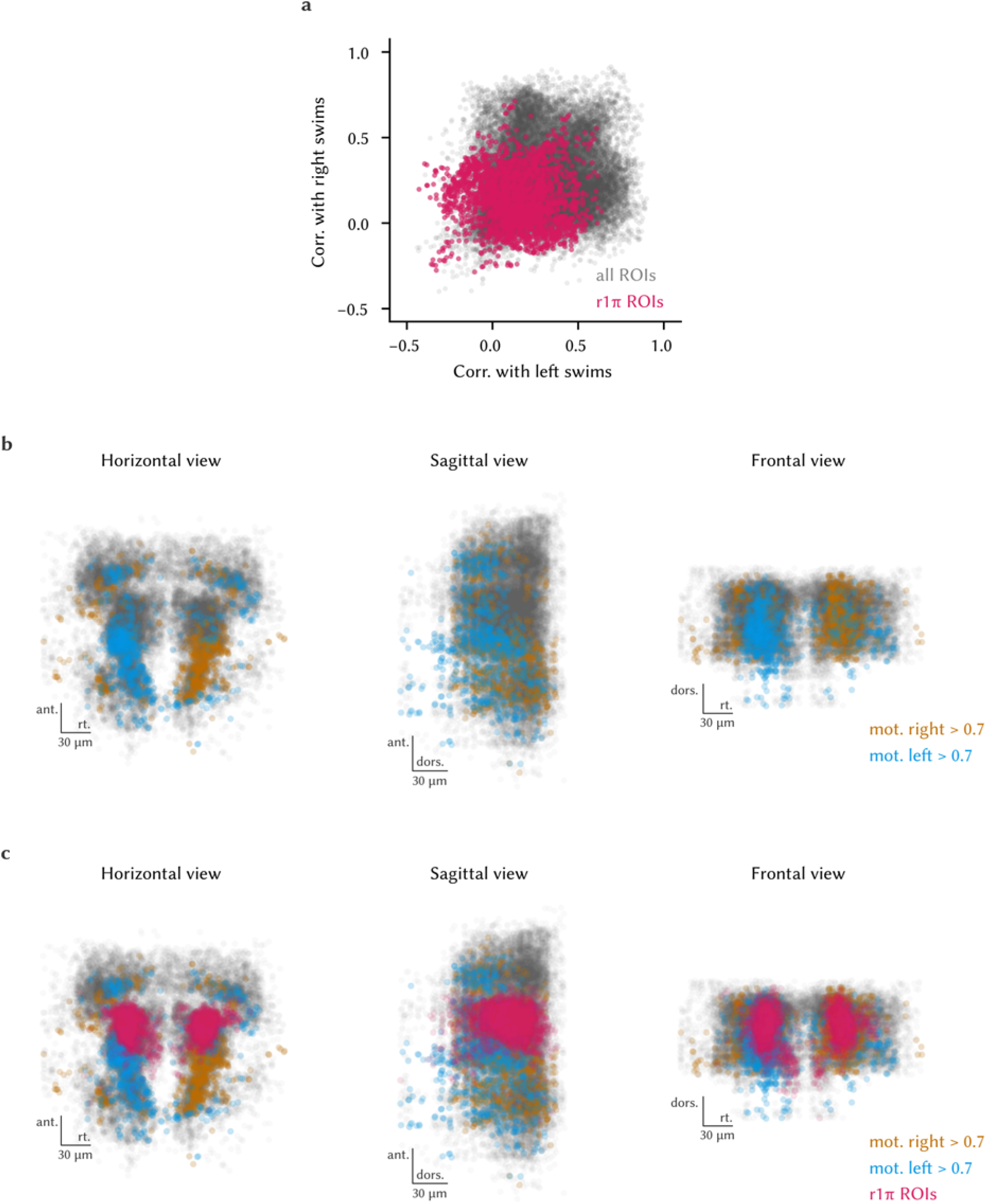
Directional-motion selective neurons in the caudal aHB. a, Correlations of all ROIs and r1π ROIs with left and right swims regressors. b, Horizontal, sagittal and frontal view showing all ROIs that have a correlation > 0.7 with a regressor for swims in one direction and < 0.7 with the regressor for swims to the opposite side (blue: left swims; golden: right swims). c, Same plot as in a, showing also all the r1π neurons.

**Supplementary Figure 11:**
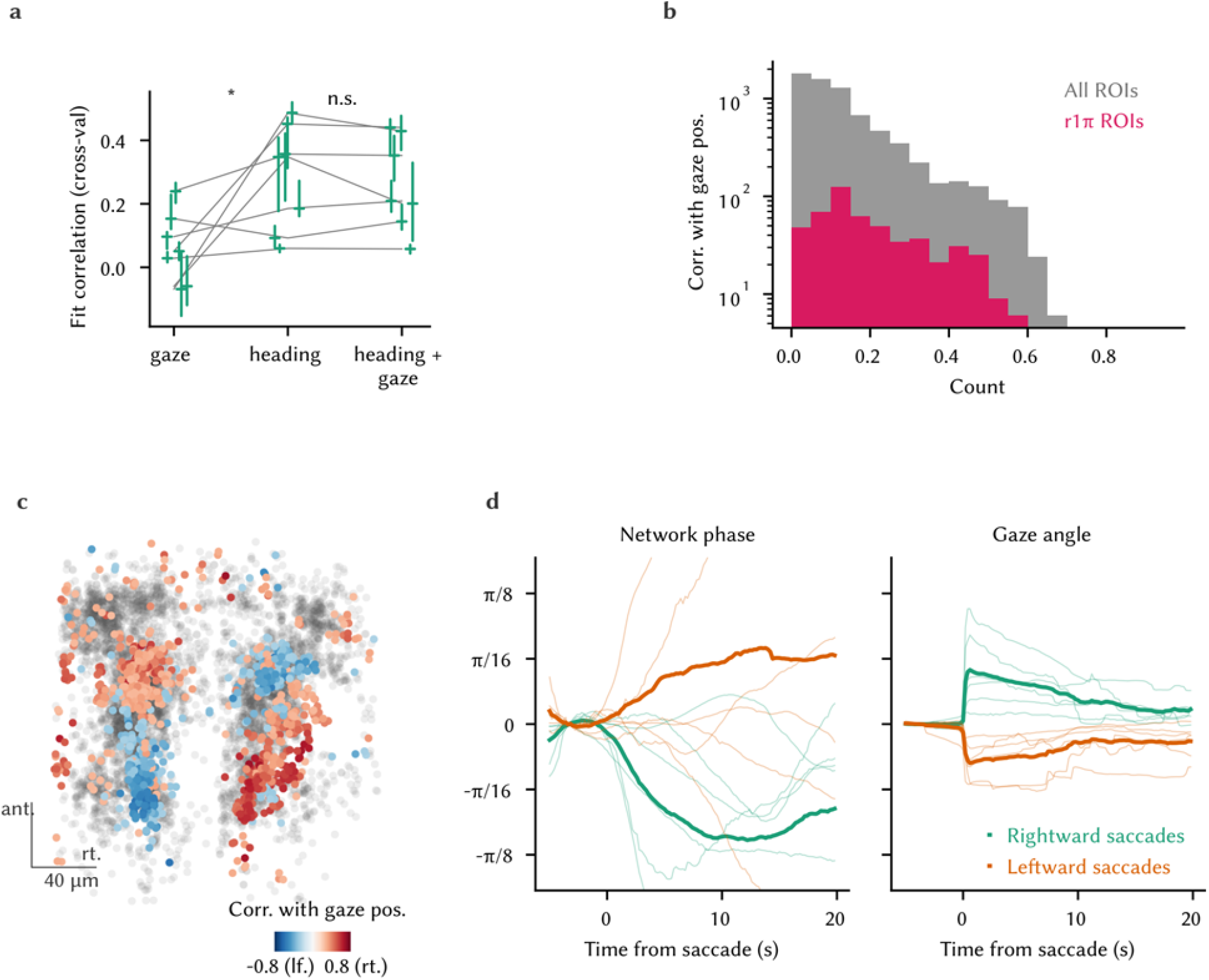
Network phase and eye motion. a, Correlation of the reconstructed test phase and the actual phase when the prediction was performed using only the gaze information, using only the heading information, or a combination of both (gaze: median = 0.0507, Q1 = −0.0155, Q3 = 0.125, n = 7 fish); (heading: median = 0.348, Q1 = 0.139, Q3 = 0.404, n = 7 fish); (gaze + heading: median = 0.209, Q1 = 0.173, Q3 = 0.391, n = 7 fish). Comparisons: Wilcoxon test, n = 7 fish. b, Histogram of the correlation of r1π neurons with a gaze position regressor, compared with the distribution obtained from all ROIs. d, Anatomical view (horizontal projection) of the correlation values of neurons with a gaze position regressor. c, *Left*, Saccade-triggered phase changes in the network, and *right*, gaze deflections for rightward (green) and leftward (orange) saccades.

**Supplementary Figure 12:**
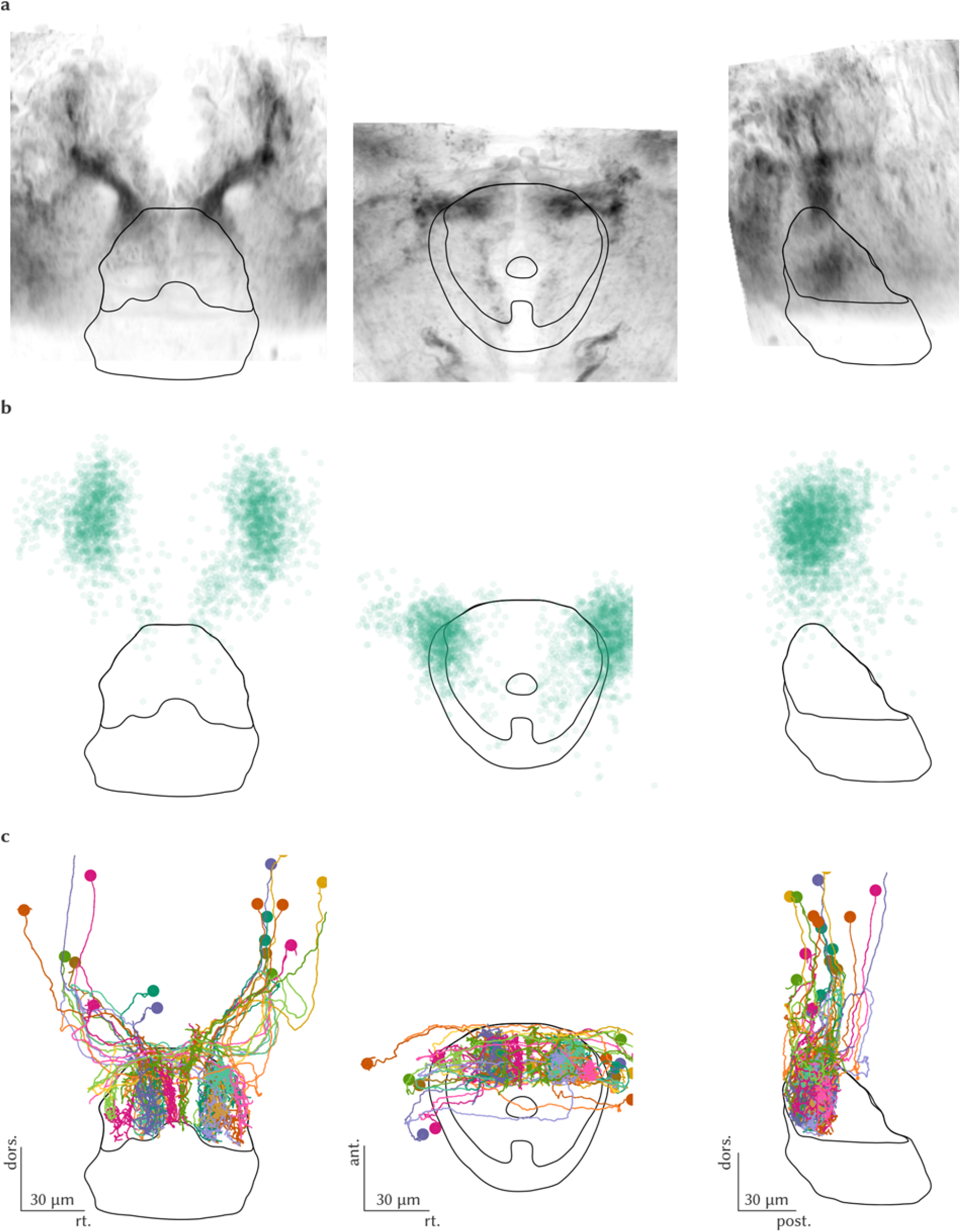
r1π neurons form reciprocal connections in the IPN. a, Frontal, horizontal and sagittal view from gad1b:Gal4, UAS:Dendra-kras stack in the region around the IPN. b, Same views, with a scatter plot representing the position of all r1π neurons from the functional dataset mapped to the IPN reference space. c, Same views, with the reconstructions from all the neurons projecting to the dIPN from the SBEM dataset, shown on their original side.

**Supplementary Figure 13:**
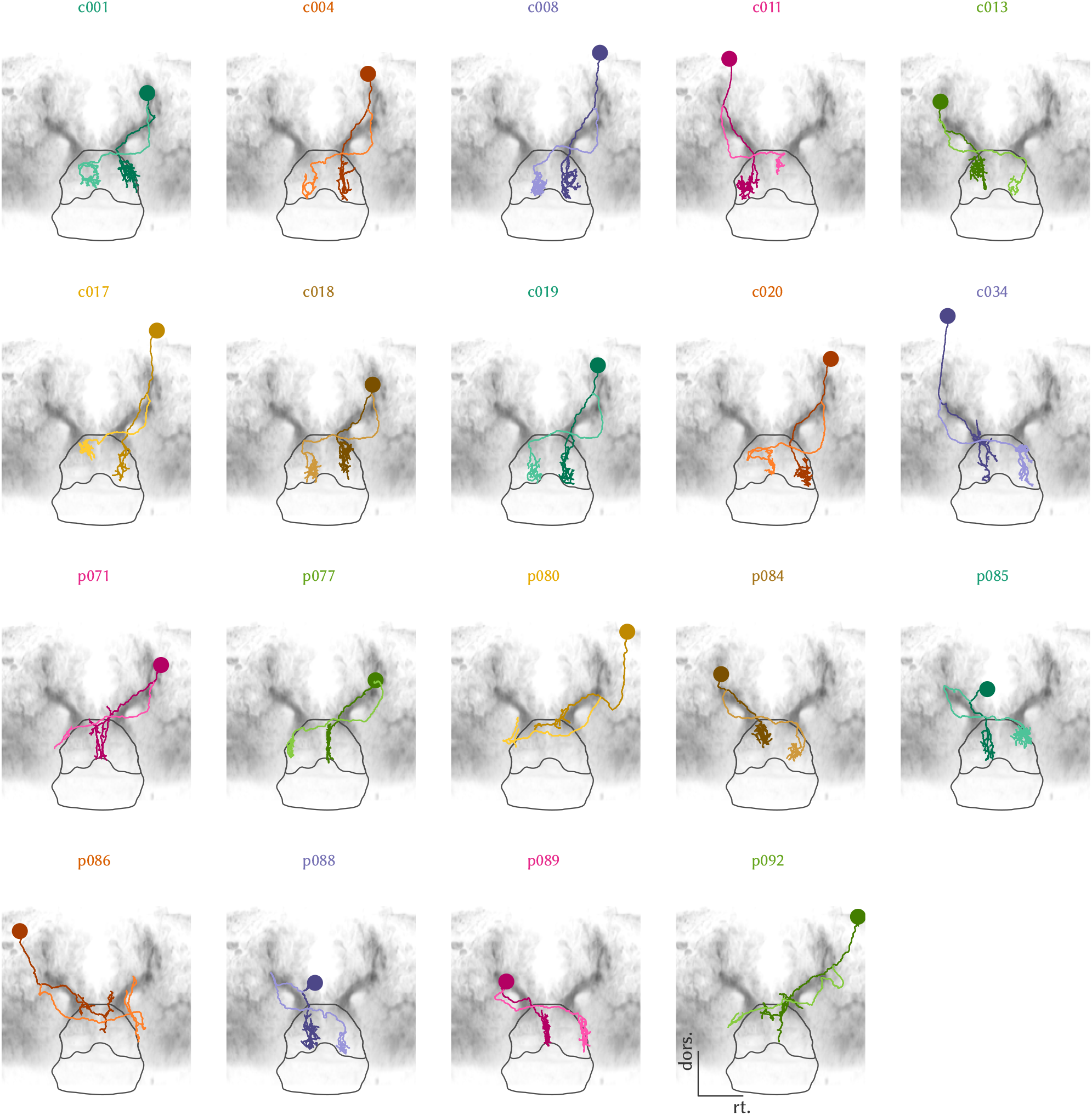
Individual plots of SBEM reconstructed neurons. Frontal view for all neurons presented in Figure 15 and Supplementary Figure 12.

**Supplementary Figure 14:**
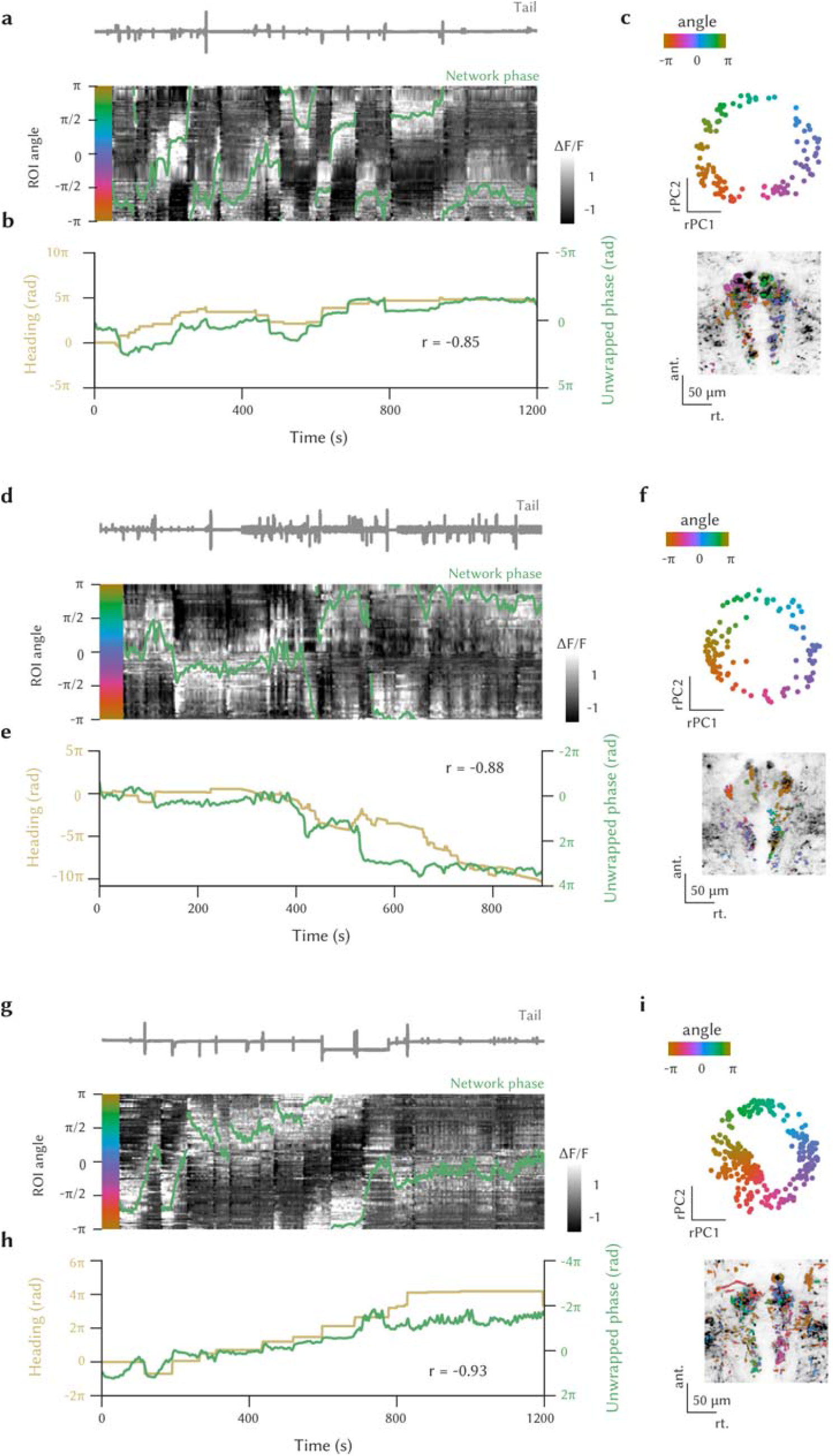
r1π neuron-like activity in the dIPN. Three example datasets are shown. a,d,g, Traces of ROIs in the dIPN showing r1π-like dynamics, sorted by angle in PC space, and phase of the network (green line). The tail trace is shown in gray on top. b,e,h, Estimated heading direction (gold) and the unwrapped network phase (green). c,f,i, *Top*, Projection over the first 2 PCs in time of all the ROI showing r1π-like activity, color-coded by angle around the circle. *Bottom*, Anatomical distribution of the same ROI, color-coded by angle in PC space. The anatomy of the recorded plane is shown in the background.

**Supplementary Figure 15:**
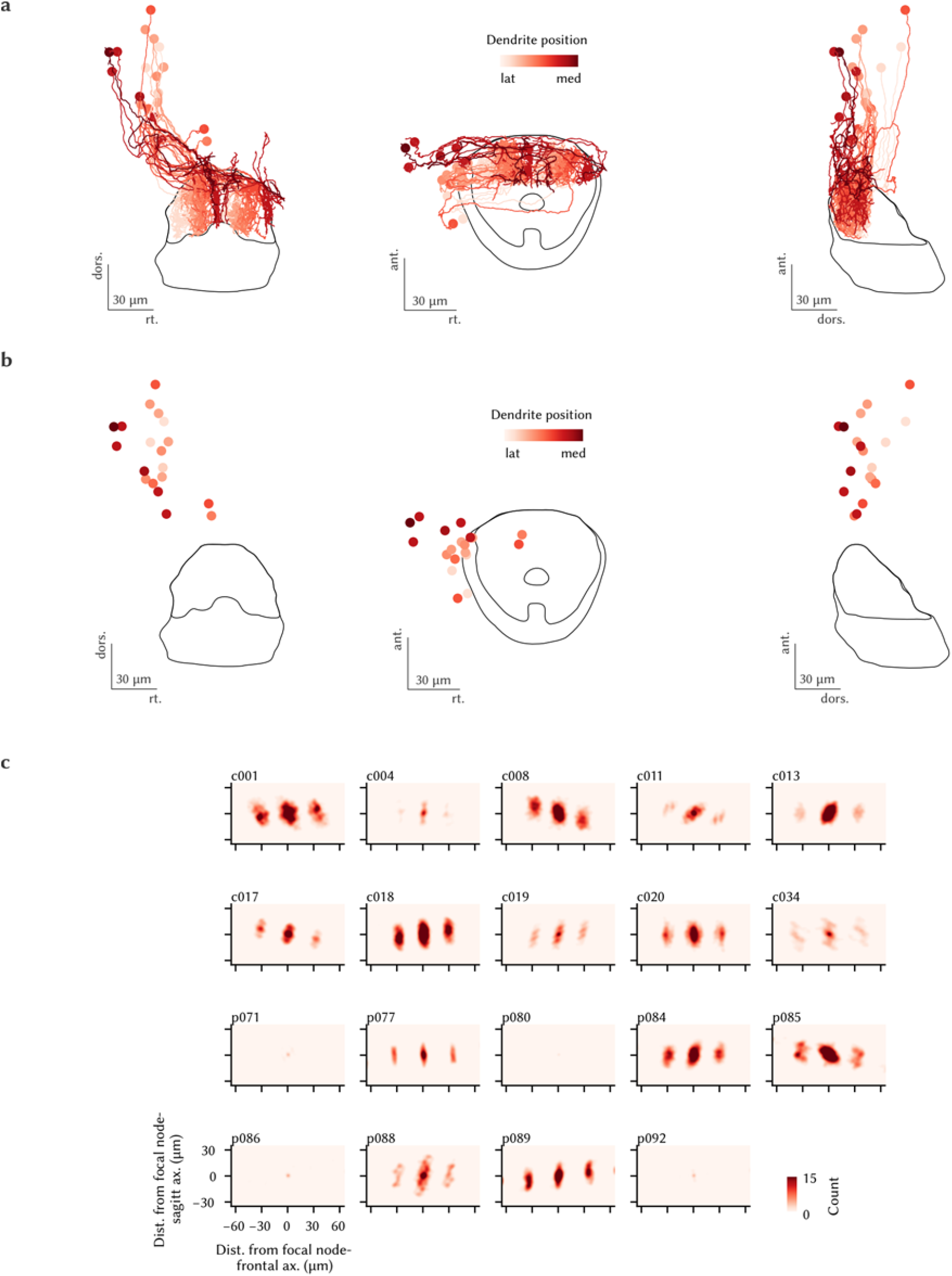
The organization of IPN projecting neurons of the aHB. a, Complete frontal, horizontal and side view for the data presented in Figure 18. b, The same views, now showing only the soma locations. c, Plots of node distances for each reconstructed neuron; those data were summed to obtain the panel in Figure 20, bottom.

**Supplementary Figure 16:**
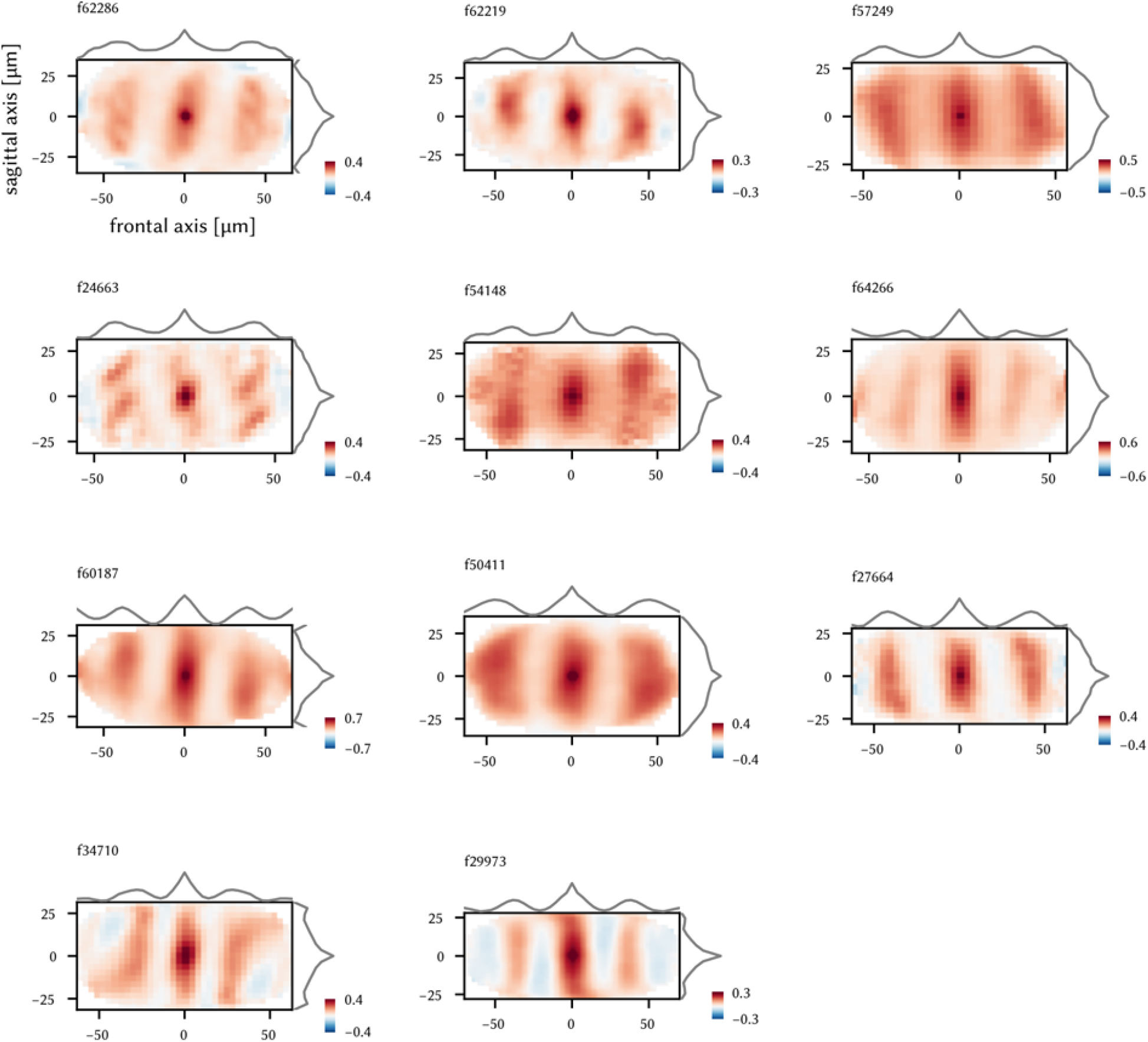
2d auto-correlation of the activity in the dIPN for all fish in the dataset. Each matrix shows the mean correlation between a focal bin and the other bins at different distances in a two-photon plane for each fish. The lines on the side show the means across each axis.

## Bibliography

Gary V. Allen and David A. Hopkins. Mamillary body in the rat: Topog-raphy and synaptology of projections from the subicular complex, prefrontal cortex, and midbrain tegmentum. The Journal of Comparative Neurology, pages 286311–336, 1989.

Isaac H. Bianco and Stephen W. Wilson. The habenular nuclei: A conserved asymmetric relay station in the vertebrate brain. Philosophical Transactions of the Royal Society B: Biological Sciences, 364(1519):1005–1020, 2009. ISSN 14712970. doi: 10.1098/rstb.2008.0213.

Rishidev Chaudhuri, Berk Gerçek, Biraj Pandey, Adrien Peyrache, and Ila Fiete. The intrinsic attractor manifold and population dynamics of a canonical cognitive circuit across waking and sleep. Nature Neuroscience, 22: 1512–1520, 2019. doi: 10.1038/s41593-019-0460-x.

Xiuye Chen and Florian Engert. Navigational strategies underlying phototaxis in larval zebrafish. Frontiers in Systems Neuroscience, 8, 2014. doi: 10.3389/fnsys.2014.00039.

Xiuye Chen, Yu Mu, Yu Hu, Aaron T. Kuan, Maxim Nikitchenko, Owen Randlett, Alex B. Chen, Jeffery P. Gavornik, Haim Sompolinsky, Florian Engert, and Misha B. Ahrens. Brain-wide organization of neuronal activity and convergent sensorimotor transformations in larval zebrafish. Neuron, 100:876–890.e5, 2018. doi: 10.1016/j.neuron.2018.09.042.

Bor Wei Cherng, Tanvir Islam, Makio Torigoe, Takashi Tsuboi, and Hitoshi Okamoto. The dorsal lateral habenula-interpeduncular nucleus pathway is essential for left-right-dependent decision making in zebrafish. Cell Reports, 32:108143, 9 2020. doi: 10.1016/j.celrep.2020.108143.

Benjamin J. Clark and Jeffrey S. Taube. Deficits in Landmark Navigation and Path Integration After Lesions of the Interpeduncular Nucleus. Behavioral Neuroscience, 123(3):490–503, 2009. ISSN 07357044. doi: 10.1037/a0015477.

Benjamin J. Clark, Asha Sarma, and Jeffrey S. Taube. Head direction cell instability in the anterior dorsal thalamus after lesions of the interpe-duncular nucleus. Journal of Neuroscience, 29(2):493–507, 2009. ISSN 02706474. doi: 10.1523/JNEUROSCI.2811-08.2009.

Antonio Contestabile and Brian A. Flumerfelt. Afferent connections of the interpeduncular nucleus and the topographic organization of the habenu lo-lnterpeduncu lar pathway: An hrp study in the rat. The Journal of Comparative Neurology, 196:253–270, 1981.

Elena I. Dragomir, Vilim Štih, and Ruben Portugues. Evidence accumulation during a sensorimotor decision task revealed by whole-brain imaging. Nature Neuroscience, 23:85–93, 2020. doi: 10.1038/s41593-019-0535-8.

Timothy W. Dunn, Yu Mu, Sujatha Narayan, Owen Randlett, Eva A. Naumann, Chao Tsung Yang, Alexander F. Schier, Jeremy Freeman, Florian Engert, and Misha B. Ahrens. Brain-wide mapping of neural activity controlling zebrafish exploratory locomotion. eLife, 5:1–29, 2016. doi: 10.7554/eLife.12741.

Yvette E. Fisher, Jenny Lu, Isabel D’Alessandro, and Rachel I. Wilson. Senso-rimotor experience remaps visual input to a heading-direction network. Nature, 576:121–125, 11 2019. doi: 10.1038/s41586-019-1772-4.

Jonathan Green, Atsuko Adachi, Kunal K. Shah, Jonathan D. Hirokawa, Pablo S. Magani, and Gaby Maimon. A neural circuit architecture for angular integration in drosophila. Nature, 546:101–106, 5 2017. doi: 10.1038/nature22343.

Henk J. Groenewegen, Sven Ahlenius, Suzanne N. Haber, Neil W. Kowall, and Walla J.H. Nauta. Cytoarchitecture, fiber connections, and some histochemical aspects of the interpeduncular nucleus in the rat. The Journal of Comparative Neurology, pages 249–65, 1986.

David Hansel and Haim Sompolinsky. Modeling feature selectivity in local cortical circuits. In Christof Koch and Idan Segev, editors, Methods in Neuronal Modeling: From Synapse to Networks, pages 499–568. The MIT Press, 1998.

Charles J. Herrick. Interpeduncular nucleus. In The Brain of the Tiger Sala-mander, pages 191–211. University of Chicago Press, 1948.

Elim Hong, Kirankumar Santhakumar, Courtney A. Akitake, Sang Jung Ahn, Christine Thisse, Bernard Thisse, Claire Wyart, Jean Marie Mangin, and Marnie E. Halpern. Cholinergic left-right asymmetry in the habenulo-interpeduncular pathway. Proceedings of the National Academy of Sciences of the United States of America, 110(52):21171–21176, 2013. ISSN 00278424. doi: 10.1073/pnas.1319566110.

Kuo Hua Huang, Misha B. Ahrens, Timothy W. Dunn, and Florian Engert. Spinal projection neurons control turning behaviors in zebrafish. Current Biology, 23:1566–1573, 8 2013. doi: 10.1016/j.cub.2013.06.044.

Brad K. Hulse and Vivek Jayaraman. Mechanisms underlying the neural computation of head direction. Annual Review of Neuroscience, 43:31–54, 7 2020. doi: 10.1146/annurev-neuro-072116-031516.

Nobuharu Iwahori, Kaori Nakamura, Sanae Kameda, and Hiroyuki Tahara. Terminal patterns of the tegmental afferents in the interpeduncular nucleus: a golgi study in the mouse. Anatomy and Embryology, 188:593–599, 1993. doi: 10.1007/BF00187015.

Kenichi Kanatani and Prasanna Rangarajan. Hyper least squares fitting of circles and ellipses. Computational Statistics and Data Analysis, 55:2197–2208, 6 2011. doi: 10.1016/j.csda.2010.12.012.

Sung Soo Kim, Hervé Rouault, Shaul Druckmann, and Vivek Jayaraman. Ring attractor dynamics in the drosophila central brain. Science, 356:849–853, 5 2017. doi: 10.1126/science.aal4835.

Rita Liu, Lisa Chang, and Gregory Wickern. The dorsal tegmental nucleus: an axoplasmic transport study. Brain Research, 310:123–132, 1984.

Cheng Lyu, L. F. Abbott, and Gaby Maimon. Building an allocentric travelling direction signal via vector computation. Nature, 601:92–97, 1 2022. doi: 10.1038/s41586-021-04067-0.

Edvard I. Moser, Emilio Kropff, and May Britt Moser. Place cells, grid cells, and the brain’s spatial representation system. Annual Review of Neuro-science, 31:69–89, 6 2008. doi: 10.1146/annurev.neuro.31.061307.090723.

Luis Puelles. Comments on the limits and internal structure of the mam-malian midbrain. Anatomy, 10:60–70, 2016. doi: 10.2399/ana.15.045.

Lely A. Quina, Julie Harris, Hongkui Zeng, and Eric E. Turner. Specific connections of the interpeduncular subnuclei reveal distinct components of the habenulopeduncular pathway. Journal of Comparative Neurology, 525(12):2632–2656, 2017. ISSN 10969861. doi: 10.1002/cne.24221.

Alexandro D. Ramirez and Emre R.F. Aksay. Ramp-to-threshold dynamics in a hindbrain population controls the timing of spontaneous saccades. Nature Communications, 12:1–19, 2021. doi: 10.1038/s41467-021-24336-w.

Johannes D. Seelig and Vivek Jayaraman. Neural dynamics for landmark orientation and angular path integration. Nature, 521:186–191, 5 2015. doi: 10.1038/nature14446.

Patricia E. Sharp, Hugh T. Blair, and Jeiwon Cho. The anatomical and computational basis of the rat head-direction cell signal. Trends in Neuro-sciences, 24:289–294, 5 2001. doi: 10.1016/S0166-2236(00)01797-5.

William E. Skaggs, James J. Knierim, Hemant S. Kudrimoti, and Bruce L. McNaughton. A model of the neural basis of the rat’s sense of direction. Advances in neural information processing systems, 7:173–180, 1995.

Marie P. Suver, Andrew M.M. Matheson, Sinekdha Sarkar, Matthew Damiata, David Schoppik, and Katherine I. Nagel. Encoding of wind direction by central neurons in drosophila. Neuron, 102:828–842.e7, 5 2019. doi: 10.1016/j.neuron.2019.03.012.

Jeffrey S. Taube. The head direction signal: Origins and sensory-motor integration. Annual Review of Neuroscience, 30:181–207, 2007. doi: 10.1146/annurev.neuro.29.051605.112854.

Jeffrey S. Taube, Robert U. Muller, and James B. Ranck. Head-direction cells recorded from the postsubiculum in freely moving rats. i. description and quantitative analysis. Journal of Neuroscience, 10:420–435, 1990. doi: 10.1523/jneurosci.10-02-00420.1990.

David Wirtshafter and Thomas R. Stratford. Evidence for gabaergic projections from the tegmental nuclei of gudden to the mammillary body in the rat. Brain Research, 630:188–194, 1993.

Sébastien Wolf, Alexis M. Dubreuil, Tommaso Bertoni, Urs Lucas Böhm, Volker Bormuth, Raphäel Candelier, Sophia Karpenko, David G. C. Hildebrand, Isaac H. Bianco, Rémi Monasson, and Georges Debrégeas. Sensorimotor computation underlying phototaxis in zebrafish. Nature Communications, 8:651, 2017. doi: 10.1038/s41467-017-00310-3.

Ryan M. Yoder and Jeffrey S. Taube. The vestibular contribution to the head direction signal and navigation. Frontiers in Integrative Neuroscience, 8, 4 2014. doi: 10.3389/fnint.2014.00032.

Kechen Zhang. Representation of spatial orientation by the intrinsic dynamics of the head-direction cell ensemble: A theory. Journal of Neuroscience, 16:2112–2126, 1996. doi: 10.1523/jneurosci.16-06-02112.1996.

## Supplementary References

Aristides B Arrenberg, Filippo Del Bene, and Herwig Baier. Optical control of zebrafish behavior with halorhodopsin. Proceedings of the National Academy of Sciences of the United States of America, 106:17968–17973, 2009. doi: 10.1073/pnas.0906252106.

Kevin L. Briggman, Moritz Helmstaedter, and Winfried Denk. Wiring specificity in the direction-selectivity circuit of the retina. Nature, 471:183–190, 2011. doi: 10.1038/nature09818.

Federico Claudi, Luigi Petrucco, Adam Tyson, Tiago Branco, Troy Margrie, and Ruben Portugues. Brainglobe atlas api: a common interface for neuroanatomical atlases. Journal of Open Source Software, 5:2668, 10 2020. doi: 10.21105/joss.02668.

Nicholas I. Fisher and Alan J. Lee. A correlation coefficient for circular data. Biometrika, 70:327–332, 1983.

Dominique Förster, Irene Arnold-Ammer, Eva Laurell, Alison J. Barker, An-tónio M. Fernandes, Karin Finger-Baier, Alessandro Filosa, Thomas O. Helmbrecht, Yvonne Kölsch, Enrico Kühn, Estuardo Robles, Krasimir Slanchev, Tod R. Thiele, Herwig Baier, and Fumi Kubo. Genetic targeting and anatomical registration of neuronal populations in the zebrafish brain with a new set of bac transgenic tools. 7:1–11, 2017. doi: 10.1038/s41598-017-04657-x.

Eleftherios Garyfallidis, Matthew Brett, Bagrat Amirbekian, Ariel Rokem, Stefan van der Walt, Maxime Descoteaux, and Ian Nimmo-Smith. Dipy, a library for the analysis of diffusion mri data. Frontiers in Neuroinformatics, 8, 2 2014. doi: 10.3389/fninf.2014.00008.

Charles R. Harris, K. Jarrod Millman, Stéfan J. van der Walt, Ralf Gommers, Pauli Virtanen, David Cournapeau, Eric Wieser, Julian Taylor, Sebastian Berg, Nathaniel J. Smith, Robert Kern, Matti Picus, Stephan Hoyer, Marten H. van Kerkwijk, Matthew Brett, Allan Haldane, Jaime Fernández del Río, Mark Wiebe, Pearu Peterson, Pierre Gérard-Marchant, Kevin Sheppard, Tyler Reddy, Warren Weckesser, Hameer Abbasi, Christoph Gohlke, and Travis E. Oliphant. Array programming with numpy. Nature, 585:357–362, 9 2020. doi: 10.1038/s41586-020-2649-2.

John D. Hunter. Matplotlib: A 2d graphics environment. Computing in Science and Engineering, 9:90–95, 2007. doi: 10.1109/MCSE.2007.55.

Eldar Insafutdinov, Leonid Pishchulin, Bjoern Andres, Mykhaylo Andriluka, and Bernt Schiele. Deepercut: A deeper, stronger and faster multi-person pose estimation model. Lecture Notes in Computer Science (including subseries Lecture Notes in Artificial Intelligence and Lecture Notes in Bioinformatics), 9906 LNCS:VII–IX, 2016. doi: 10.1007/978-3-319-46466-4.

Michael Kunst, Eva Laurell, Nouwar Mokayes, Anna Kramer, Fumi Kubo, António M Fernandes, Dominique Förster, Marco Dal Maschio, and Herwig Baier. A cellular-resolution atlas of the larval zebrafish brain. Neuron, 103(1):21–38, 2019.

James A. Lister, Christie P. Robertson, Thierry Lepage, Stephen L. Johnson, and David W. Raible. nacre encodes a zebrafish microphthalmia-related protein that regulatesneural-crest-derived pigment cell fate. Development, 126:3757–3767, 1999. doi: https://doi.org/10.1242/dev.126.17.3757.

Alexander Mathis, Pranav Mamidanna, Kevin M. Cury, Taiga Abe, Venkatesh N. Murthy, Mackenzie Weygandt Mathis, and Matthias Bethge. Deeplabcut: markerless pose estimation of user-defined body parts with deep learning. Nature Neuroscience, 21:1281–1289, 9 2018. doi: 10.1038/s41593-018-0209-y.

Tanmay Nath, Alexander Mathis, An Chi Chen, Amir Patel, Matthias Bethge, and Mackenzie Weygandt Mathis. Using deeplabcut for 3d markerless pose estimation across species and behaviors. Nature Protocols, 14: 2152–2176, 7 2019. doi: 10.1038/s41596-019-0176-0.

Huy Bang Nguyen, Truc Quynh Thai, Sei Saitoh, Bao Wu, Yurika Saitoh, Satoshi Shimo, Hiroshi Fujitani, Hirohide Otobe, and Nobuhiko Ohno. Conductive resins improve charging and resolution of acquired images in electron microscopic volume imaging. Scientific Reports, 6:1–10, 2016. doi: 10.1038/srep23721.

Marius Pachitariu, Carsen Stringer, Mario Dipoppa, Sylvia Schröder, L. Federico Rossi, Henry Dalgleish, Matteo Carandini, and Kenneth Harris. Suite2p: beyond 10,000 neurons with standard two-photon microscopy. bioRxiv, page 061507, 2016. doi: 10.1101/061507.

Fabian Pedregosa, Ron Weiss, Matthieu Brucher, GaÃńl Varoquaux, Alexandre Gramfort, Vincent Michel, Bertrand Thirion, Olivier Grisel, Mathieu Blondel, Peter Prettenhofer, Ron Weiss, Vincent Dubourg, Jake Vanderplas, Alexandre Passos, David Cournapeau, Matthieu Brucher, Matthieu Perrot, and édouard Duchesnay. Scikit-learn: Machine learning in python. Journal of Machine Learning Research, 12:2825–2830, 2011.

Torsten Rohlfing and Calvin R. Maurer. Nonrigid image registration in shared-memory multiprocessor environments with application to brains, breasts, and bees. IEEE Transactions on Information Technology in Biomedicine, 7:16–25, 3 2003. doi: 10.1109/TITB.2003.808506.

Vilim Štih, Luigi Petrucco, Andreas M. Kist, and Ruben Portugues. Stytra: An open-source, integrated system for stimulation, tracking and closed-loop behavioral experiments. PLoS Computational Biology, 15, 2019. doi: 10.1371/journal.pcbi.1006699.

Vilim Štih, You Kure Wu, Emanuele Paoli, Diego Asua, and Ruben Portugues. portugueslab/brunoise: Alpha, October 2020. URL https://doi.org/10.5281/zenodo.4122064.

Vilim Štih, Diego Asua, Luigi Petrucco, Federico Puppo, and Ruben Portugues. Sashimi, January 2022a. URL https://doi.org/10.5281/zenodo.5932227.

Vilim Štih, Luigi Petrucco, Ot Prat, Hagar Lavian, and Ruben Portugues. Bouter, January 2022b. URL https://doi.org/10.5281/zenodo.5931684.

Fabian N. Svara, Jörgen Kornfeld, Winfried Denk, and Johann H. Bollmann. Volume em reconstruction of spinal cord reveals wiring specificity in speed-related motor circuits. Cell Reports, 23:2942–2954, 2018. doi: 10.1016/j.celrep.2018.05.023.

Michael A. Taylor, Gilles C. Vanwalleghem, Itia A. Favre-Bulle, and Ethan K. Scott. Diffuse light-sheet microscopy for stripe-free calcium imaging of neural populations. Journal of biophotonics, 11(12):e201800088, 2018.

Tod R. Thiele, Joseph C. Donovan, and Herwig Baier. Descending control of swim posture by a midbrain nucleus in zebrafish. Neuron, 83:679–691, 8 2014. doi: 10.1016/j.neuron.2014.04.018.

Pauli Virtanen, Ralf Gommers, Travis E. Oliphant, Matt Haberland, Tyler Reddy, David Cournapeau, Evgeni Burovski, Pearu Peterson, Warren Weckesser, Jonathan Bright, Stéfan J. van der Walt, Matthew Brett, Joshua Wilson, K. Jarrod Millman, Nikolay Mayorov, Andrew R.J. Nelson, Eric Jones, Robert Kern, Eric Larson, C. J. Carey, İlhan Polat, Yu Feng, Eric W. Moore, Jake VanderPlas, Denis Laxalde, Josef Perktold, Robert Cimrman, Ian Henriksen, E. A. Quintero, Charles R. Harris, Anne M. Archibald, Antônio H. Ribeiro, Fabian Pedregosa, Paul van Mulbregt, Aditya Vijaykumar, Alessandro Pietro Bardelli, Alex Rothberg, Andreas Hilboll, Andreas Kloeckner, Anthony Scopatz, Antony Lee, Ariel Rokem, C. Nathan Woods, Chad Fulton, Charles Masson, Christian Häggström, Clark Fitzgerald, David A. Nicholson, David R. Hagen, Dmitrii V. Pasechnik, Emanuele Olivetti, Eric Martin, Eric Wieser, Fabrice Silva, Felix Lenders, Florian Wilhelm, G. Young, Gavin A. Price, Gert Ludwig Ingold, Gregory E. Allen, Gregory R. Lee, Hervé Audren, Irvin Probst, Jörg P. Dietrich, Jacob Silterra, James T. Webber, Janko Slavič, Joel Nothman, Johannes Buchner, Johannes Kulick, Johannes L. Schönberger, José Vinícius de Miranda Cardoso, Joscha Reimer, Joseph Harrington, Juan Luis Cano Rodríguez, Juan Nunez-Iglesias, Justin Kuczynski, Kevin Tritz, Martin Thoma, Matthew Newville, Matthias Kümmerer, Maximilian Bolingbroke, Michael Tartre, Mikhail Pak, Nathaniel J. Smith, Nikolai Nowaczyk, Nikolay Shebanov, Oleksandr Pavlyk, Per A. Brodtkorb, Perry Lee, Robert T. McGibbon, Roman Feldbauer, Sam Lewis, Sam Tygier, Scott Sievert, Sebastiano Vigna, Stefan Peterson, Surhud More, Tadeusz Pudlik, Takuya Oshima, Thomas J. Pingel, Thomas P. Robitaille, Thomas Spura, Thouis R. Jones, Tim Cera, Tim Leslie, Tiziano Zito, Tom Krauss, Utkarsh Upadhyay, Yaroslav O. Halchenko, and Yoshiki Vázquez-Baeza. Scipy 1.0: fundamental algorithms for scientific computing in python. Nature Methods, 17:261–272, 3 2020. doi: 10.1038/s41592-019-0686-2.

